# *Zymoseptoria tritici* effectors structurally related to killer proteins UmV-KP4 and UmV-KP6 are toxic to fungi, and define extended protein families in fungi

**DOI:** 10.1101/2024.10.14.618152

**Authors:** Karine de Guillen, Léa Mammri, Jérôme Gracy, André Padilla, Philippe Barthe, François Hoh, Mounia Lahfa, Justine Rouffet, Yohann Petit-Houdenot, Thomas Kroj, Marc-Henri Lebrun

## Abstract

Fungal effectors play crucial roles in plant infection. Despite low sequence identity, effectors were recently classified into structural families. In this study, we have elucidated the structures of Zt-NIP1 and Mycgr3-91409 effectors of the wheat fungal pathogen *Zymoseptoria tritici,* using X-ray crystallography and NMR. These effectors displayed a structural homology with, respectively, KP4 and KP6α killer toxins, from UmV dsRNA viruses infecting the corn fungal pathogen *Ustilago maydis*. Consequently, Zt-NIP1 and Mycgr3-91409 were renamed Zt-KP4-1 and Zt-KP6-1. Orthologs and paralogs of Zt-KP4-1 and Zt-KP6-1 were identified in *Zymoseptoria*, but not in other fungi, except Ecp2 effectors related to Zt-KP4-1. Assessment of the biological activities of Zt-KP6-1 and Zt-KP4-1 revealed their toxicity to fungi such as *Botrytis cinerea* and *Z. tritici*, but not to wheat. A novel pipeline relying on Foldseek and cysteine-pattern constrained HMM searches of AlphaFold2 predicted structures from Uniprot generated a comprehensive inventory of KP4 and KP6 proteins in fungi and plants. A structure-based classification of these proteins revealed four KP4 and three KP6 structural super families. This classification provided far-reaching hypotheses on their biological function and evolution. This unifying structural framework highlights the power of structure determination for the classification of effectors, and their functional investigation.

## INTRODUCTION

Fungal plant pathogens secrete large repertoires of effectors during host plant infection mostly composed of small proteins without enzymatic functions [1,2]. Most fungal effectors display no or very limited protein sequence similarities [3,4]. However, the comparison of their three-dimensional structures allows their classification into families of structurally related proteins [3]. This strategy was successful in identifying a large number of avirulence (AVR) and ToxB-like effectors from the fungal rice pathogen *Pyricularia oryzae* sharing the same conserved fold and called MAX effectors [5–7]. The RNase-like proteins associated with haustoria (RALPH) effectors of the cereal powdery mildew fungus *Blumeria graminis* were also grouped into a family of structurally related proteins using modeling and experimental structure determination [8–11]. Additional structural families of fungal effectors were identified using experimental structure determination and modeling in *Leptosphaeria maculans* [12–15], *Fusarium oxysporum* [16,17], and *Venturia inaequalis* [18]. Recently, the systematic prediction of structures of secreted fungal proteins with different programs including AlphaFold2 (AF2) [19], revealed novel structural families of fungal effectors [20,21].

To identify novel families of fungal effectors, we elucidated the three-dimensional structures of the secreted proteins Zt-Mycgr3-91409 and Zt-NIP1 from the wheat fungal pathogen *Zymoseptoria tritici* by X-ray diffraction (XRD) and nuclear magnetic resonance (NMR). Zt-Mycgr3-91409 is an effector upregulated during infection of wheat leaves [22] previously suggested to be a MAX effector [5]. Zt-NIP1 is expressed during infection of wheat [22,23] and induces mild chlorotic lesions on wheat leaves [24]. Unexpectedly, the structures of Zt-Mycgr3-91409 and Zt-NIP1 shared structural similarities with UmV-KP6α and UmV-KP4 killer proteins (KP) toxins encoded by dsRNA viruses infecting the corn fungal pathogen *Ustilago maydis*. UmV-KP6α and UmV-KP4 kill susceptible *U. maydis* strains, while UmV infected strains are resistant to the killer toxin they produce [25,26]. Consequently, we renamed Zt-Mycgr3-91409 to Zt-KP6-1 and Zt-NIP1 to Zt-KP4-1. Bioassays revealed that Zt-KP6-1 and Zt-KP4-1 were toxic to fungi, but not to wheat. Proteins structurally related to Zt-KP6-1 or Zt-KP4-1 were widely distributed across fungi, and classified into a unifying framework of structural families.

## RESULTS

### Zt-Mycgr3-91409-2 is a structural homolog of UmV-KP6 killer toxin

The *Zt-Mycgr3-91409* gene model was curated according to transcript evidence [27] (Sup Figure 1A). The corresponding protein, Zt-Mycgr3-91409-2 (SMR48303, A0A2H1G421) was expressed without its secretion signal in *Escherichia coli* and purified by affinity and size exclusion chromatography (SEC) (Sup File S1, Sup Figure 2). High-resolution diffracting crystals were obtained after optimization of Li_2_SO_4_ concentration and pH and soaked in tantalum. Analyses of X-Rays Diffraction data (XRD) at 1.36 Å resolution showed that crystals belonged to the *P*2_1_2_1_2_1_ space group with two copies of the protein per asymmetric unit. The structure of Zt-Mycgr3-91409-2 was solved by single-wavelength anomalous diffraction (SAD), revealing a protein with a typical α/β sandwich structure (Figure 1, Sup Table 1). This fold has two α-helices and four β-strands arranged in an antiparallel β-sheet with one hairpin connection formed by β2-β3 and two βαβ connections, which placed the α-helices in an antiparallel manner on one side of the β-sheet.

**Figure 1.**
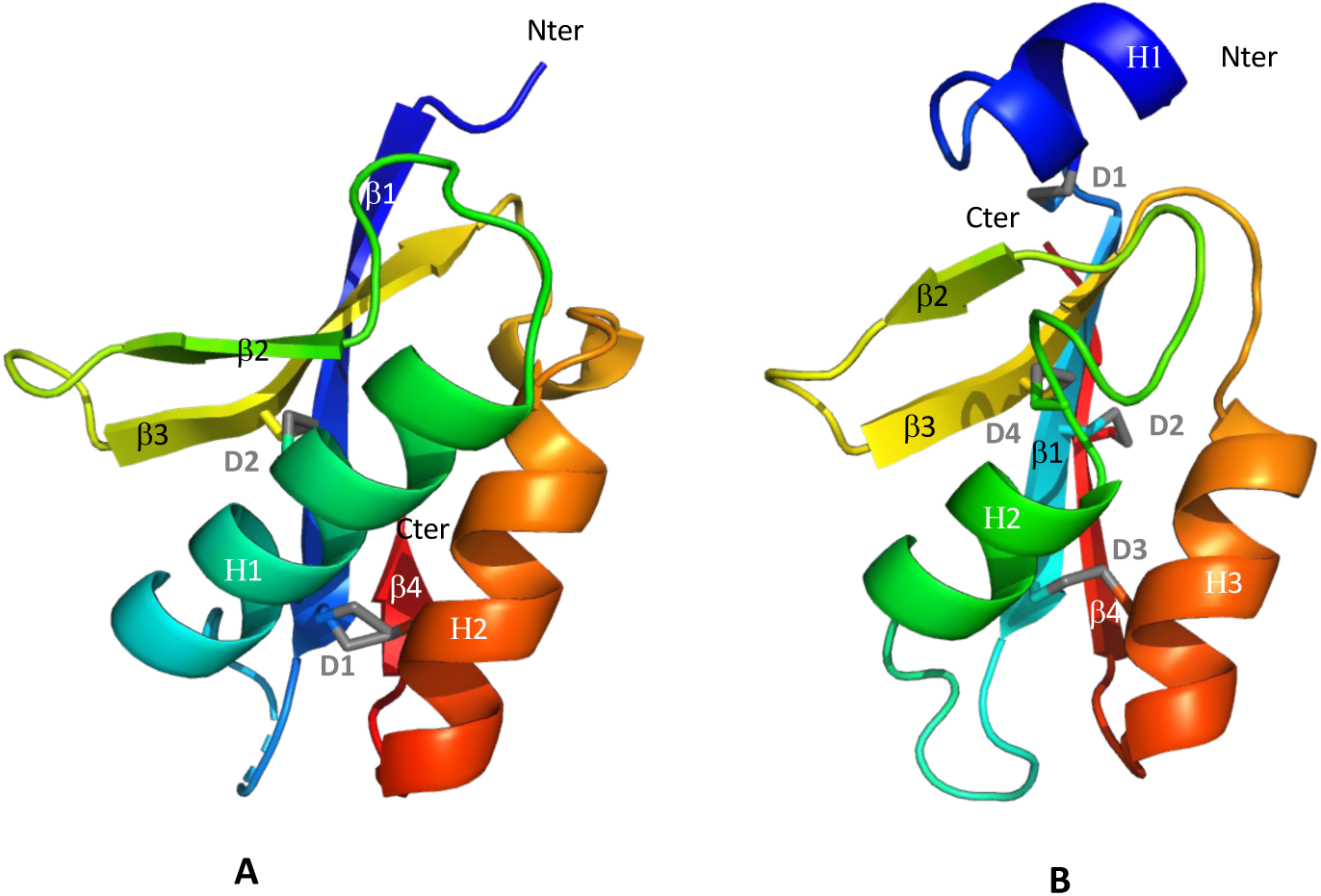
Crystal structures of (A) Zt-KP6-1 (6QPK chain A) and (B) UmV-KP6α (4GVB chain A). (A) Zt-Mycgr3-91409-2 was renamed to Zt-KP6-1 according to its structural homology to UmV-KP6α (see text). Structures were coloured in rainbow colours from blue (N terminus) to red (C terminus). For Zt-KP6-1, the loop connecting β1 (in blue) to H1 (in cyan) was not observed in the density map and is shown as a blue dashed line. Disulphide bonds are shown in grey and labelled Dx. For the D1 (C26-C86) bond in Zt-KP6-1, both conformations are shown.

The α-helix H2 was preceded by a 3^10^ helix formed by residues 77 to 89 (numbering according to the mature protein). Both α-helices (H1 and H2) were tightly packed against the β-sheet, with one disulphide bond formed between cysteine residues in β1 (Cys26) and H2 (Cys86) and another disulphide bond formed between cysteine residues in H1 (Cys43) and β3 (Cys65), making the fold very compact. Alternative conformations were observed for Cys26 involving the disulphide bond with Cys86. The interface area between the two monomers observed in the crystal, determined by PDBePISA, was estimated to be 435.8Å^2^. The Complexation Significance Score (CSS) was 0, indicating that the dimer has no biological relevance and resulted from crystal packing. NMR studies of the protein labelled with ^15^N revealed a high similarity to the crystal structure, with an r.m.s.d. of 0.7 ± 0.6 Å on structured parts. However, the protein was not dimeric in solution, as shown by the narrowness of the ^1^H-^15^N HSQC spectrum peaks (Sup Figure 3 and Sup Table 2), and the single elution profile at 95 min observed in SEC corresponding to a retention time of a 9 kDa globular protein (Sup Figure 2).

A DALI analysis [28] revealed a structural similarity between Zt-Mycgr3-91409-2 and UmV-KP6α (PDB 4GVB, Figure 1, Z-score of 6.7 and r.m.s.d of 2.5 Å for 63 aligned residues). UmV-KP6α is produced together with UmV-KP6β by cleavage of the UmV-KP6 pre-protein encoded by a dsRNA virus infecting *U. maydis,* a fungal pathogen causing corn smut disease [26]. Both mature proteins have a highly similar fold [26,29] defined by two α-helices (H1 and H2) positioned on one side of a 4-stranded β-sheet, with a β4-β1-H1-β3-H2-β2 topology (KP6 fold), also observed for Zt-KP6-1 (Sup Figure 4). All three proteins have two right handed β-α-β split crossovers, which distinguish the KP6 fold from toxins of similar size [30]. Another characteristic of the KP6 fold was the two disulphide bonds that linked the antiparallel α-helices to the central β-sheet of the protein (Sup. Figure. 4). However, Zt-KP6-1 did not have the N-terminal hydrophobic α-helix extension found in UmV-KP6α. According to these structural similarities, Zt-Mycgr3-91409-2 was renamed Zt-KP6-1.

### Zt-NIP1 is a structural homolog of *UmV*-KP4 killer toxin

Zt-NIP1 was identified as a wheat necrosis-inducing protein produced by IPO323 *Z. tritici* isolate during wheat leaf infection [24]. However, the CDS at the *Zt-NIP1* locus in IPO323 genome (*Zt-Mycgr3-106176*) encoded a protein with a sequence unrelated to NIP1. We re-annotated the locus by predicting a new CDS in agreement with transcript evidence and almost identical to NIP1 (99% identity) [27] (Sup Figure 1B). Zt-NIP1 (SMR6055, A0A2H1H404) was expressed in *E. coli* (Sup File S1) and purified by affinity chromatography and SEC. SEC revealed an equilibrium between Zt-NIP1 monomers and dimers in solution (Sup Figure 5). High-resolution diffracting crystals were obtained and belonged to the *H3* space group with 8 copies of the protein per asymmetric unit (Sup Table 1). The Zt-NIP1 structure was solved by molecular replacement using an AF2 model (Figure 2) revealing a typical α/β sandwich fold with a single split βαβ motif and two disulphide bonds. A main antiparallel 5-stranded β-sheet formed by β1-β3-β4-β6-β7, was packed against two α-helices (H1 and H2) and combined with a second, small antiparallel two-stranded β-sheet comprising β2 and β5 (with 4 residues each). The fold was held together by two disulphide bonds formed between cysteine residues in β2 (Cys39) and β7 (Cys142), and H1 (Cys60) and β4 (Cys89). Chains E-H, G-B, F-C, and D-A formed dimers, through the main 5-stranded β-sheet with an average interface area of 852.2 Å^2^ determined by PDBePISA. These dimers exhibited a CSS score of 1.0, supporting the biological significance of dimer formation (Sup Figure 6).

**Figure 2.**
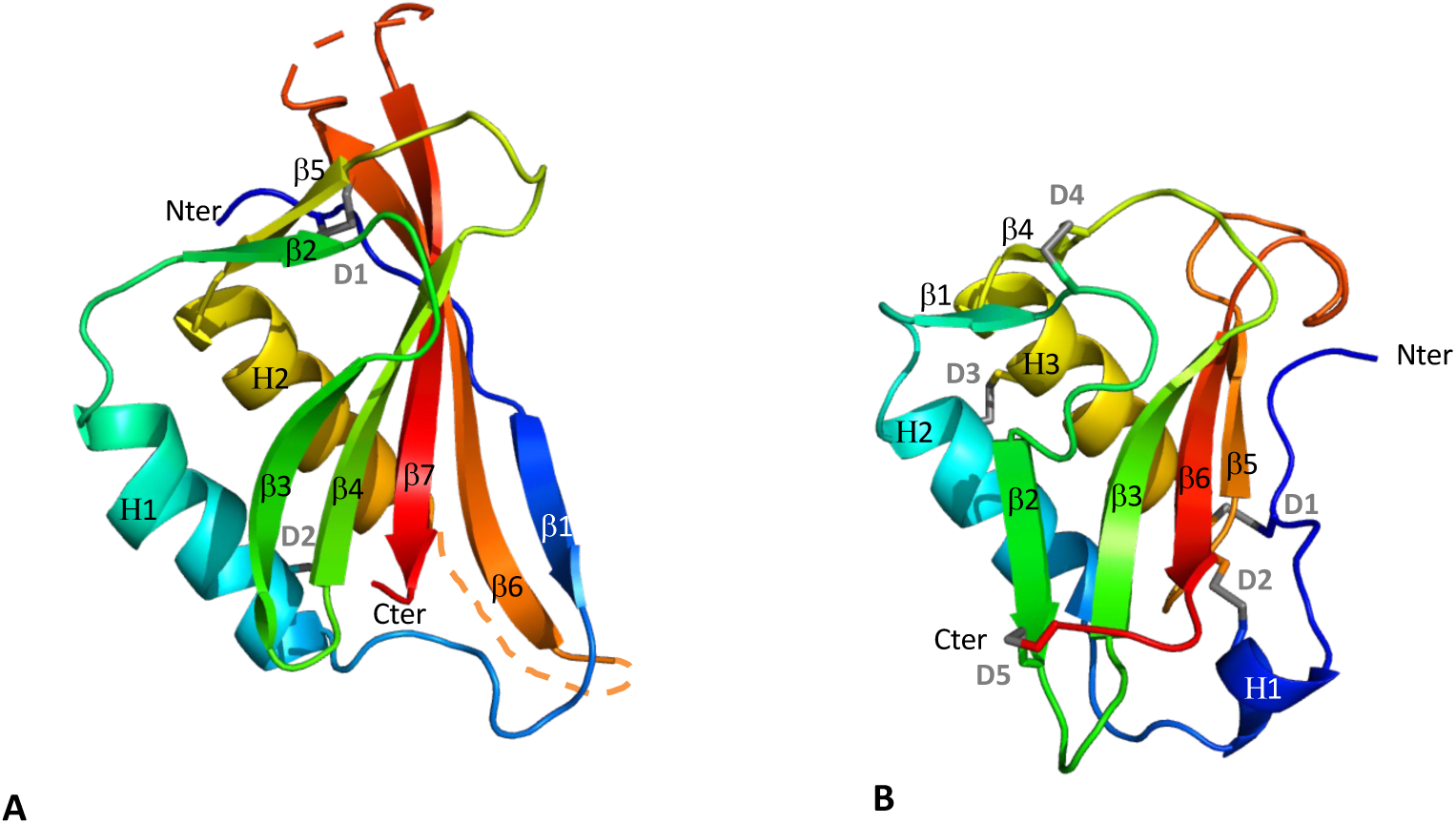
Crystal structures of (A) Zt-KP4-1 (8ACX chain A) and (B) UmV-KP4 (1KPT chain A). (A) Zt-NIP1 was renamed to Zt-KP4-1 according to its structural similarity to UmV-KP4 using DALI (see main text). Structures were coloured in rainbow colours from blue (N terminus) to red (C terminus). For Zt-KP4-1 the loops connecting β6 to β7 and β6 to H1 were not observed in the density map and were shown as dashed lines. Disulphide bonds are shown in grey and labelled Dx.

A DALI analysis [28], revealed structural similarity between Zt-NIP1 and UmV-KP4 (PDB 1KPT, Figure 2). UmV-KP4, like UmV-KP6 is encoded by a dsRNA virus infecting *U. maydis* [26]. The two proteins superposed with a Z-score of 7.3 and a r.m.s.d value of 2.4 Å (82 residues aligned). UmV-KP4 [31,32] belongs to the α/β sandwich family of proteins and is composed of seven β strands and three α-helices. Similar to Zt-NIP1, UmV-KP4 has two helices (H2 and H3) in a left-handed βαβ crossover conformation, which is a rare structural feature (Sup Figure 7) [30]. The main difference between the two proteins is the presence of a β-strand (β1) at the N-terminus for Zt-NIP1 instead of a short α-helix (H1) for UmV-KP4. The two proteins also differ in the number and location of disulphide bonds. While Zt-NIP1 has two disulphide bonds (as described above), UmV-KP4 has five. Based on the structural similarity of these proteins, Zt-NIP1 was renamed to Zt-KP4-1.

### Zt-KP4 and Zt-KP6 are toxic to fungi

When Zt-KP4-1 or Zt-KP6-1 (2 µM) were infiltrated into leaves of the wheat cultivar Taichung 29, which is susceptible to *Z. tritici*, they did not induce visible signs of toxicity such as leaf chlorosis or necrosis. The necrotrophic effector ToxA from *P. nodorum* [33], used as a positive control in this experiment, induced typical necrotic lesions. To test if Zt-KP4-1 and Zt-KP6-1 were toxic to fungi, we monitored their effect on the growth of the model fungal plant pathogen *Botrytis cinerea* and of *Z. tritici* isolate IPO323, which carries both Zt-KP4-1 and Zt-KP6-1. The protein concentrations used in these experiments were within the range of 0.5 to 5.5 µM. These concentrations were below those commonly used for the investigation of fungal toxic proteins which are generally in the range of 20 µM [34,35]. Zt-KP4-1 strongly inhibited the growth of both fungi at 5.5 µM (91-95 % inhibition, Table 1). At 1.65 µM, it only inhibited *Z. tritici* (48 % inhibition, Table 1). Zt-KP6-1 strongly inhibited the growth of *B. cinerea* (94 % inhibition at 5.5 µM, Table 1), but had only a weaker effect on *Z. tritici* growth (24 % inhibition at 5.5 µM, Table 1). These results showed that Zt-KP4-1 and Zt-KP6-1 are toxic to fungi, but not to wheat. While Zt-KP4-1 displayed the same toxicity on *Z. tritici* and *B. cinerea,* Zt-KP6-1 showed selectivity, as it was four times more active on *B. cinerea* than on *Z. tritici*.

**Table 1.** Toxicity of Zt-KP4-1 and Zt-KP6-1 to fungi. The percentages of *B. cinerea* and *Z. tritici* growth inhibition were calculated for different concentrations of ToxA, Zt-KP4-1and Zt-KP6-1. Values are the mean ± Standard Error of the Mean (SEM) from three experiments.

|  |  | % inhibition |  |
| --- | --- | --- | --- |
|  |  | *Botrytis cinerea* | *Zymoseptoria tritici* |
| Protein concentration (μM) |  |  |  |
| TOX A | 5.50 | 25 ± 1.4 | 0 |
|  | 1.65 | 0 | 0 |
|  | 0.55 | 0 | 0 |
| Zt-KP4-1 | 5.50 | 91 ± 3.2 | 95 ± 1.4 |
|  | 1.65 | 0 | 48 ± 2.2 |
|  | 0.55 | 0 | 25 ± 1.4 |
| Zt-KP6-1 | 5.50 | 94 ± 2.0 | 24 ± 1.2 |
|  | 1.65 | 0 | 0 |
|  | 0.55 | 0 | 0 |

### Identification of Zt-KP6-1 homologs in *Zymoseptoria*

Proteins with sequences highly similar to Zt-KP6-1 were identified in other strains of *Z. tritici* [36] and *Z. pseudotritici* (Zp-KP6-1, 92% identity, Figure 3), but not in other fungi, including available genomes from other *Zymoseptoria* species. Proteins with a low sequence identity to Zt-KP6-1 (28-30%) were identified in *Z. tritici* (Zt-KP6-2) and other *Zymoseptoria* species (KP6-2 from *Z. brevis, Z. arbibiliaie* and *Z. pseudotritici*, Figure 3) but not in other fungi. The structure of the Zt-KP6-2 protein predicted by AF2 was highly similar to that of Zt-KP6-1 (Z-score of 12.1; r.m.s.d of 1.9 Å for 73 aligned residues, Sup Figure 8). However, it contained an 11 amino acid C-terminal sequence with two cysteine residues conserved in all KP6-2 proteins, but absent from KP6-1 proteins. The phylogenetic analysis of KP6-1 and KP6-2 proteins highlighted two clades gathering orthologous proteins from different *Zymoseptoria* species and suggesting that *Zymoseptoria* KP6-1 and KP6-2 are paralogs originating from the duplication of an ancestral gene before the divergence of the respective species (Figure 3).

**Figure 3.**
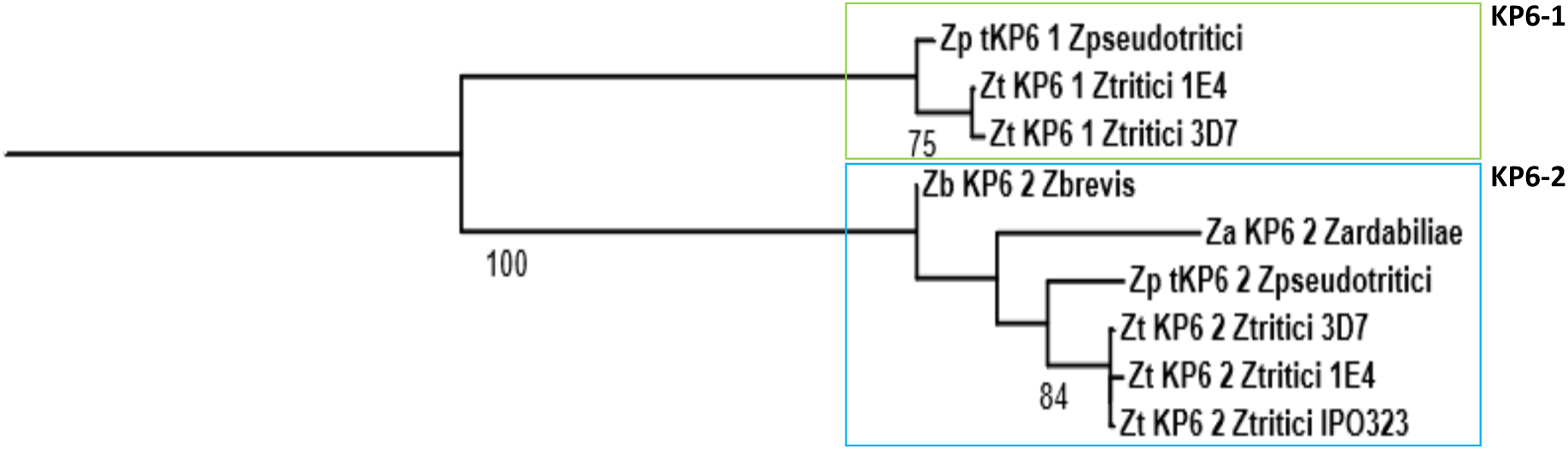
Phylogenetic tree of *Zymoseptoria* KP6-1 and KP6-2. MAFFT alignment, BMGE cleaning, PhyML Phylogenetic tree using NGPhylogeny.fr. [37] Bootstrap values > 70% are indicated under each branch after the supported node. Midpoint tree rooting. Zt-KP6-1_Ztritici_3D7: Uniprot A0A1X7RM13 (identical to Zt-KP6-1_IPO323); Zt-KP6-1_Ztritici_1E4: Uniprot A0A2H1G421, PDB 6QKP; Zp-KP6-1_Zpseudotritici: JGI Zymps1_802032; Zt-KP6-2_Ztritici_1E4: Uniprot A0A2H1H0K4; Zt-KP6-2_Ztritici_IPO323: Uniprot F9XLY9; Zt-KP6-2_Ztritici_3D7: Uniprot A0A1X7S5L2; Zb-KP6-2_Zbrevis: Uniprot A0A0F4GMG5; Za-KP6-2_Zardabiliae: JGI Za_774343 ; Zp-KP6-2_Zpseudotritici: JGI Zymps1_805921.

Using a novel pipeline based on FoldSeek and Hmmer (FHAT, <u>F</u>oldseek <u>H</u>mmer <u>A</u>lphafold <u>T</u>Malign, Sup Figure 9), we identified four additional candidate KP6 proteins in *Zymoseptoria.* Visual inspection of their AF2-predicted structures revealed important overlap with the Zt-KP6-1 structure for three of them: Zt-KP6-4, Zt-KP6-5 and Zb-KP6-6. The remaining one, Zt-KP6-3, displayed a low structural similarity with KP6-1 (DALI score of 0) and it was excluded from further analyses. Overall, *Zymoseptoria* had five different KP6 proteins with low or no sequence similarity, but sharing a similar three-dimensional structure.

### Identification of Zt-KP4-1 homologs in *Zymoseptoria*

Proteins with sequences highly similar to Zt-KP4-1 were identified in other strains of *Z. tritici* [36] (Figure 4), but not in other *Zymoseptoria* species. Proteins with a low sequence identity to Zt-KP4-1 (33-35%) were identified in *Z. tritici* (Zt-KP4-2; Zt-KP4-3) and *Z. brevis* (Zb-KP4-2; Zb-KP4-3). The AF2 structures of the Zt-KP4-2 and Zt-KP4-3 proteins predicted by AF2 were highly similar to that of Zt-KP4-1 (Zt-KP4-2: Z-score of 16.5; r.m.s.d of 1.6 Å for 111 aligned residues, Zt-KP4-3: Z-score of 15.8; r.m.s.d of 1.7 Å for 110 aligned residues Sup Figure 10). BlastP searches of the proteomes of other fungi with Zt-KP4-1 or Zt-KP4-2 as queries, retrieved proteins related to the effector ECP2 (ExtraCellular Protein 2) from the tomato pathogen *Passalora fulva* (syn. *Cladosporium fulvum*) [38,39] Yet, the similarity scores were low (28-32% identity, e-value 10^-20^). A phylogenetic analysis of *Zymoseptoria* KP4-1, KP4-2 and KP4-3 proteins including ECP2 proteins as outgroups [38].The *Zymoseptoria* KP4-1, KP4-2 and KP4-3 proteins clustered as three independent clades gathering orthologous proteins from *Zymoseptoria* rooted by the ECP2 proteins (Figure 4). This topology suggested that KP4-1, KP4-2 and KP4-3 originated from duplications of an ancestral ECP2-related gene, prior to the divergence of *Zymoseptoria* species.

**Figure 4.**
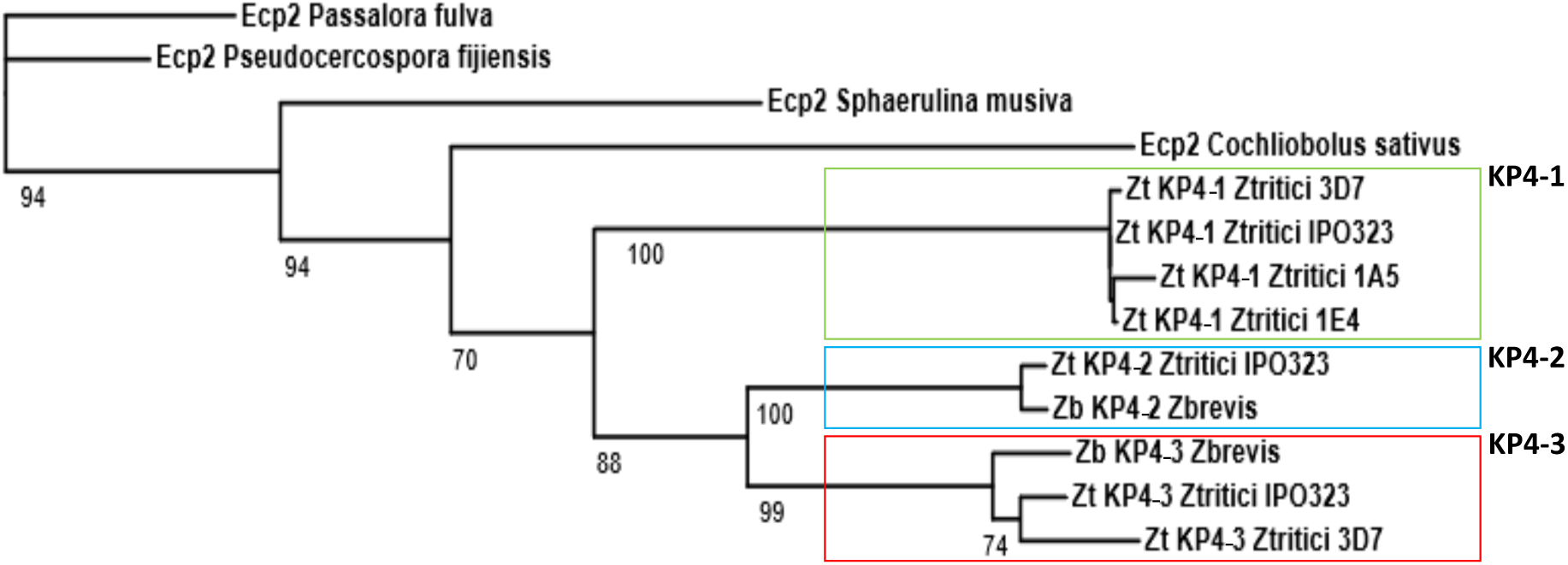
Phylogenetic tree of *Zymoseptoria* KP4-1, KP4-2, and KP4-3. MAFFT alignment, BMGE cleaning, PhyML Phylogenetic tree using NGPhylogeny.fr [37]. Bootstrap values > 70% are indicated under each branch after the supported node. Tree root with Ecp2_Passalora fulva and Ecp2_Pseudocercospora fijiensis as outgroups. Zt-KP4-1_Ztritici_IPO323: re-annotated NIP1/Mycgr3-106176 see Sup. Fig1; Zt-KP4-1_Ztritici_1E4: Uniprot A0A2H1H404; Zt-KP4-1_Ztritici_3D7: Uniprot A0A1X7S6U3; Zt-KP4-1_Ztritici_1A5: Uniprot A0A1Y6M012; Zt-KP4-2_Ztritici_IPO323: Mycgr3-107904, Uniprot F9X223; Zb-KP4-2_Zbrevis: KJY02238, Uniprot A0A0F4GYX0; Zt-KP4-3_Ztritici_IPO323: Mycgr3-111636, Uniprot F9XQ02; Zt-KP4-3_Ztritici_3D7: Uniprot A0A1X7S9R6; Zb-KP4-3_Zbrevis: KJX96086, Uniprot A0A0F4GF74; Ecp2_Passalora fulva: Uniprot Q00365; Ecp2_Pseudocercospora fijiensis: Uniprot M2Z6V3; Ecp2_Sphaerulina musiva: Uniprot N1QK49; Ecp2_Cochliobolus sativus: Uniprot M2TE34

A previous large-scale comparative genomics study of fungal effectors outlined the HCE2 superfamily widely distributed in fungi (HCE2: Homologs of *C. fulvum* Ecp2)[39]. This analysis established a novel pfam domain (HCE2, PF14856) to identify proteins related to Ecp2. We detected the HCE2 domain in Zt-KP4-1, Zt-KP4-2 and Zt-KP4-3, which is consistent with their sequence similarity to *C. fulvum* Ecp2. However, we did not detect additional *Z. tritici* proteins with a HCE2 domain. Using FHAT (Sup Figure 9), we identified six additional candidate KP4 proteins in *Zymoseptoria*. Visual inspection of their AF2-predicted structures revealed important overlap with the Zt-KP4-1 structure for four of them: Zt-KP4-5, Zt-KP4-6, Zt-KP4-7 and Zt-KP4-9. The remaining ones, Zt-KP4-4 and Zt-KP4-8, displayed only a low structural similarity to Zt-KP4-1 despite the presence of the same number of α−helices and β strands and a similar organization. In Zt-KP4-4, the structures of the β2 and β5 strands were not well defined (twisted), and the β2-β3 and β5-β6 interactions were reduced. In Zt-KP4-8, the interactions between the β1 and β4 strands were missing. These two proteins were therefore excluded from further analyses. Overall, *Zymoseptoria* has seven KP4 proteins with low or no sequence similarity, but sharing the same three-dimensional structure.

### *Zymoseptoria KP4* and *KP6* are expressed during infection

Published IPO323 RNA-Seq data [22] showed that *Zt-KP4-1* was the most highly expressed *Zt-KP4* gene during infection of wheat leaves (25 x beta-tubulin *TUB1*), with maximum expression at 9 dpi (Sup Fig. 11). It was also expressed during *in vitro* growth, although to a lesser extent. *Zt-KP4-3* was highly expressed during infection at 9 and 14 dpi (10 x *TUB1*), and during *in vitro* growth. *Zt-KP4-2* was constitutively expressed at a level similar to *TUB1*. *Zt-KP4-5, Zt-KP4-6* and *Zt-KP4-7* were expressed at low levels (1/10^th^ of *TUB1*) in all analysed conditions, while *Zt-KP4-9* was not expressed. *Zt-KP6-1*, *Zt-KP6-2*, *Zt-KP6-4* and *Zt-KP6-6* displayed a similar expression pattern with maximum expression during infection at 9 dpi (1 to 2 x *TUB1*), and highly reduced expression during *in vitro* growth (100 fold less than during infection). *Zt-KP6-5* was not expressed. In summary, this survey showed that most *Zymoseptoria KP4* and *KP6* were over-expressed during infection at the switch from the asymptomatic phase to the necrotrophic phase (9-14 dpi). This switch is associated with the massive colonization of the apoplast of the infected wheat leaf by infectious hyphae of *Z. tritici*, and the invasion of sub-stomatal cavities where they differentiate into pycnidia [40].

### Proteins with a KP6 fold are widespread in fungi

To establish a comprehensive inventory of proteins structurally related to Zt-KP6-1 in fungi, we searched 1014 fungal genomes for structurally related proteins using a novel Fold-Seek/Hmmer pipeline (Sup Figure 9). After filtering of results for significant structural similarity to Zt-KP6-1 or UmV-KP6α using TM align (TM-score above 0.5) [41], this analysis identified 2556 KP6 candidates belonging to 81 AFDB50 clusters (kclXX-Y), which we merged into 17 meta-clusters (kclXX) (Sup Figure 9). Visual inspection of the structures of one representative protein from each meta-cluster rejected 10 meta-clusters with topologies different from the typical KP6 fold. The remaining 7 KP6 FHAT meta-clusters gathered 1307 proteins (final KP6 FHAT database) including, Zt-KP6-1 (kcl06), Zt-KP6-2 (kcl06), Zt-KP6-4 (kcl02), Zt-KP6-5 (kcl07), Zt-KP6 (kcl07) (sup Table 3). Several fungal effectors have been described as structurally related to UmV-KP6α: ECP28 from *C. fulvum* [42], AvrLm10A and AvrLm6 from *L. maculans* [14,15], SIX5 from *Fusarium oxysporum* [14], BAS4 from *Pyricularia oryzae* [43], and Cb-Nip1 from *Cercospora beticola* [44]. In addition, the systematic analysis of effector structures of *V. inaequalis* and *L. maculans* found 108 and 13 proteins structurally related to KP6, respectively [15,18]. Only few of these proteins were identified in our analysis, since most sequences were not available in Uniprot. A visual inspection of the structural overlaps between these additional KP6 proteins and Zt-KP6-1 rejected only Cb-NIP1, since its AF2 model was not sufficiently structured to allow its superposition to Zt-KP6-1. To search for structural relationships among KP6 proteins, we selected at least one representative of each of the 7 KP6 FHAT meta-clusters, one representative of each KP6 family identified in *L. maculans* (8 families) [15], one representative of each major KP6 family identified in *V. inaequalis* (4 families) [18] as well as UmV-KP6α and UmV-KP6β. This selection of 35 KP6 included all *Zymoseptoria* KP6 (Zt-KP6-1, 2, 4, 5, 6) and known KP6 (Six5, AvrLm10A, BAS4, AvrLm6, ECP28). The similarity between the structures of these 35 proteins was assessed using a principal component analysis (PCA) with DALI Z-scores. This analysis revealed three major structural super families (Groups 1, 2 and 3: Figure 5). Group-1 included AvrLm6 (kcl04-C) and AvrLm10A (kcl01-L) from *L. maculans* and SIX5 (kcl01-L) from *F. oxysporum,* but no KP6 proteins from *Zymoseptoria.* Group-2 contained Zt-KP6-4 (kcl02-D), ECP28-1/2 (kcl14-A), ECP28-3 (kcl14-B), BAS4 (kcl02-G), representatives of kcl01-C, kcl04-A, and kcl11-A, as well as UmV-KP6β. Group-3 was composed of Zt-KP6-1 (kcl06-B), UmV-KP6α, Zt-KP6-2 (kcl06-A), Zt-KP6-6 (kcl07-A), and a representative of kcl02-A. Taken together, this large-scale comparative structural analysis provided a first comprehensive description of the fungal KP6 effectors. In addition, it provided a reliable classification of the KP6 proteins into three major super families (groups in Figure 5) for follow-up studies.

**Figure 5.**
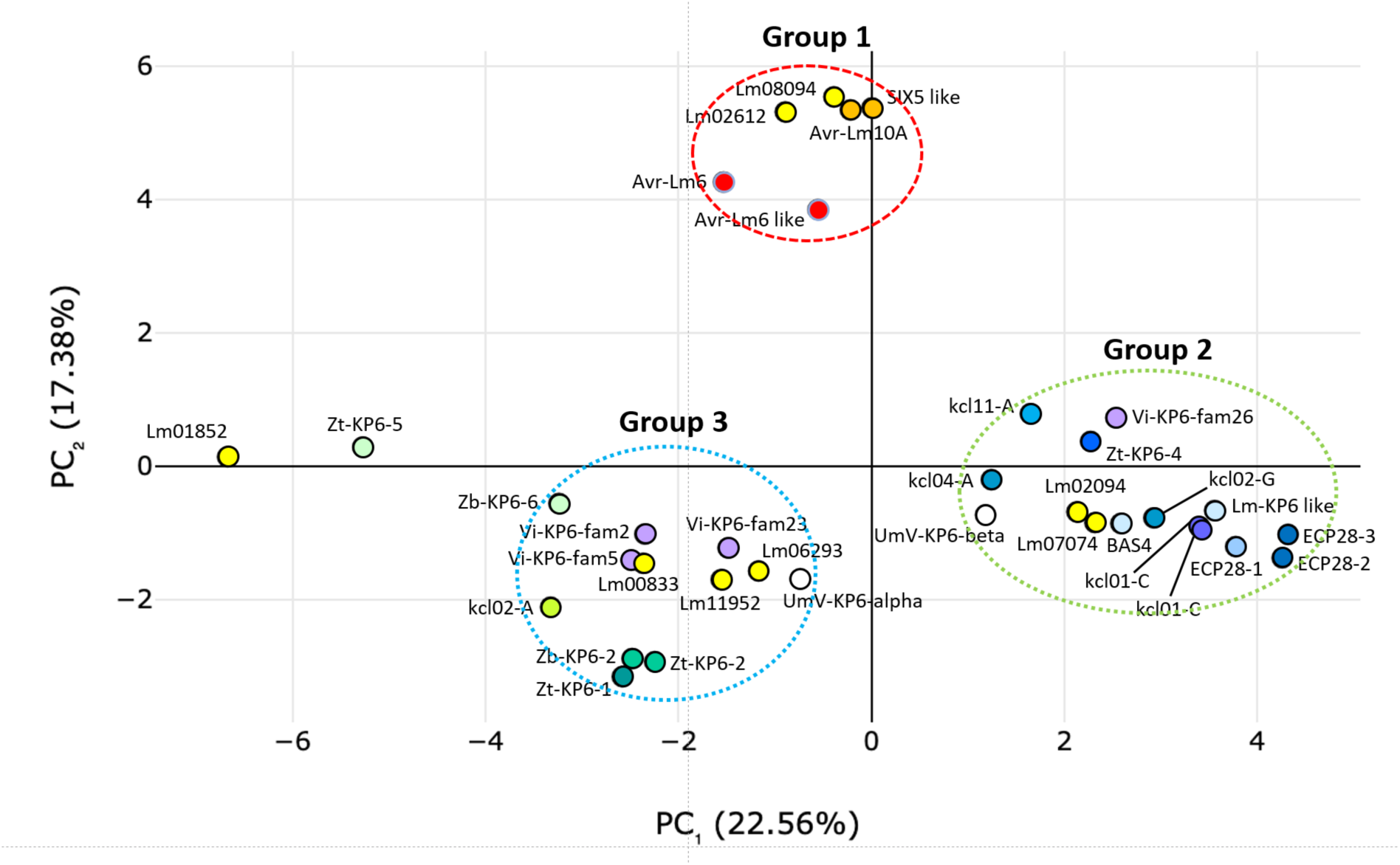
Classification of fungal KP6 families according to their structures. DALI all by all analysis followed by a PCA using a DALI score matrix [28]. meta-cluster: kclXX, AFDB50 cluster: kclXX-Y, (number of proteins in the AFDB50 cluster), Uniprot number of the representative. kcl01-C, (100), A0A4V3HRP1, M1W1H7; kcl01-L, (65), SIX5 = A0A010RIE2, AvrLm10A = E5A720_LEPMJ; kcl02-A, (5), A0A4R8TM13; kcl02-D, (138), Zt-KP6-4 = A0A2H1GZA7; kcl02-G, (321), A0A135SXI2; BAS4 = BAS4_MAGO7, Lm-KP6 = E5A0W1_LEPMJ; kcl04-A, (1), F9FAC2; kcl04-C, (25), AvrLm6 = A0A010QJV2; AvrLm6 = E5R515_LEPMJ; kcl06-A, (6), Zb-KP6-2 = A0A0F4GMG5, Zt-KP6-2 = A0A2H1H0K4; kcl06-B, (3), Zt-KP6-1 = A0A2H1G421/6QKP; kcl07-A, (5), Zt-KP6-5 = A0A2H1FMK1, Zb-KP6-6 = A0A0F4GGQ1; kcl11-A, (6), A0A317AKI4; kcl14-A, (93), ECP28-2 = A0A1P8YXM1, ECP28-3 = A0A1P8YXM5; kcl14-B, (14), ECP28-1 = A0A1P8YXQ5. LmXXXX, Leptosphaeria KP6 proteins, (8 families), [15]; Vi-KP6-famXX, Venturia KP6 proteins, (4 families), [18]; White circle, UmV-KP6α, 4gvbA, White circle, UmV-KP6β, 4gvbB

### KP4 proteins are abundant and widely distributed in fungi

In addition to the HCE2 domain (PF14856) found in Zt-KP4-1, Zt-KP4-2 and Zt-KP4-3 [39], the pfam database comprises a KP4 domain (PF09044) initially defined by the analysis of the *F. graminearum* proteins Fg-KP4-1, Fg-KP4-2 and Fg-KP4-3 related to UmV-KP4. While the HCE2 domain relied on a pattern dominated by three conserved cysteine and a W/F/Y motif, the KP4 domain was defined by a pattern involving four conserved cysteine, three conserved glycine and two W/F/Y motifs [45,46]. Interestingly, we found no overlap in proteins retrieved from Uniprot with either the HCE2 or the KP4 pfam domain. While the HCE2 domain was identified, as expected, in Zt-KP4-1, Zt-KP4-2 and Zt-KP4-3, the KP4 domain was not identified in a *Zymoseptoria* protein. To establish a comprehensive inventory of the KP4 proteins, we searched 1014 fungal genomes with FHAT using Zt-KP4-1 and UmV-KP4 as references (Sup Figure 9). We identified 4381 KP4 candidates grouped into 153 AFDB50 clusters that we merged into 44 FHAT meta-clusters. Visual inspection of the structures of one representative protein from each meta-cluster rejected 31 meta-clusters, which did not have the typical KP4 fold. The remaining 13 FHAT meta-clusters gathered 3249 proteins with a KP4 fold (final KP4 FHAT database) including Ecp2 (kcl01), Zt-KP4-1 (kcl01), Zt-KP4-2 (kcl07), Zt-KP4-3 (kcl07), Zt-KP4-5 (kcl18), Zt-KP4-6 (kcl09), Zt-KP4-7 (kcl09) and Zt-KP4-9 (kcl02) (sup Table 4). To search for structural relationships among KP4 proteins, we selected at least one representative of each of the 13 KP4 FHAT meta-clusters. This selection included known KP4 proteins such as Fg-KP4-1 and Fg-KP4-2 which defined the KP4 pfam domain [45,46] and one Ecp2 (Paf-ECP2) used for defining the HCE2 pfam domain [39]. Additional known KP4 proteins such as Fg-KP4-3 [46], UmV-KP4 and two Ecp2 proteins (Sm-ECP2, Psf-ECP2) used for the phylogenetic analysis of Zt-KP4-1 (Figure 4) were also selected. This selection of 25 KP4 included all *Zymoseptoria* KP4 (Zt-KP4-1, 2, 3, 5, 6, 7, 9). The similarity between the structures of these 25 proteins was assessed using a principal component analysis (PCA) with DALI Z-scores. This analysis revealed four major structural super families (Groups 1, 2, 3, 4: Figure 6). Group-1 was composed of Fg-KP4-1, Fg-KP4-2 and Fg-KP4-3, UmV-KP4 and the representative of kcl02-A. Group-2 contained Zt-KP4-1, Zt-KP4-2, Zt-KP4-3 and Ecp2 proteins, as well as the kcl07-A, kcl10-A and kcl14-A representatives. Group-3 included the representatives of kcl08-A and kcl38-A. Group-4 contained all other KP4 proteins, including Zt-KP4-5, Zt-KP4-6, Zt-KP4-7 and Zt-KP4-9. Interestingly, all Group-1 proteins had a pfam KP4 domain, while all proteins from Group-2 had an HCE2 domain. Proteins from Group-3 and Group-4 did not have Pfam domains, with the exception of the representatives of kcl38-A (PF15474) and kcl21-A (PF21691). This genome wide comparative structural analysis provided a novel comprehensive inventory of plant and fungal KP4 proteins that include the Ecp2 superfamilly. Their structure-based classification in four super families (groups in Figure 6) clarified the relationship between proteins with either a HCE2 or a KP4 pfam domain providing a novel framework for this large family of effectors.

**Figure 6.**
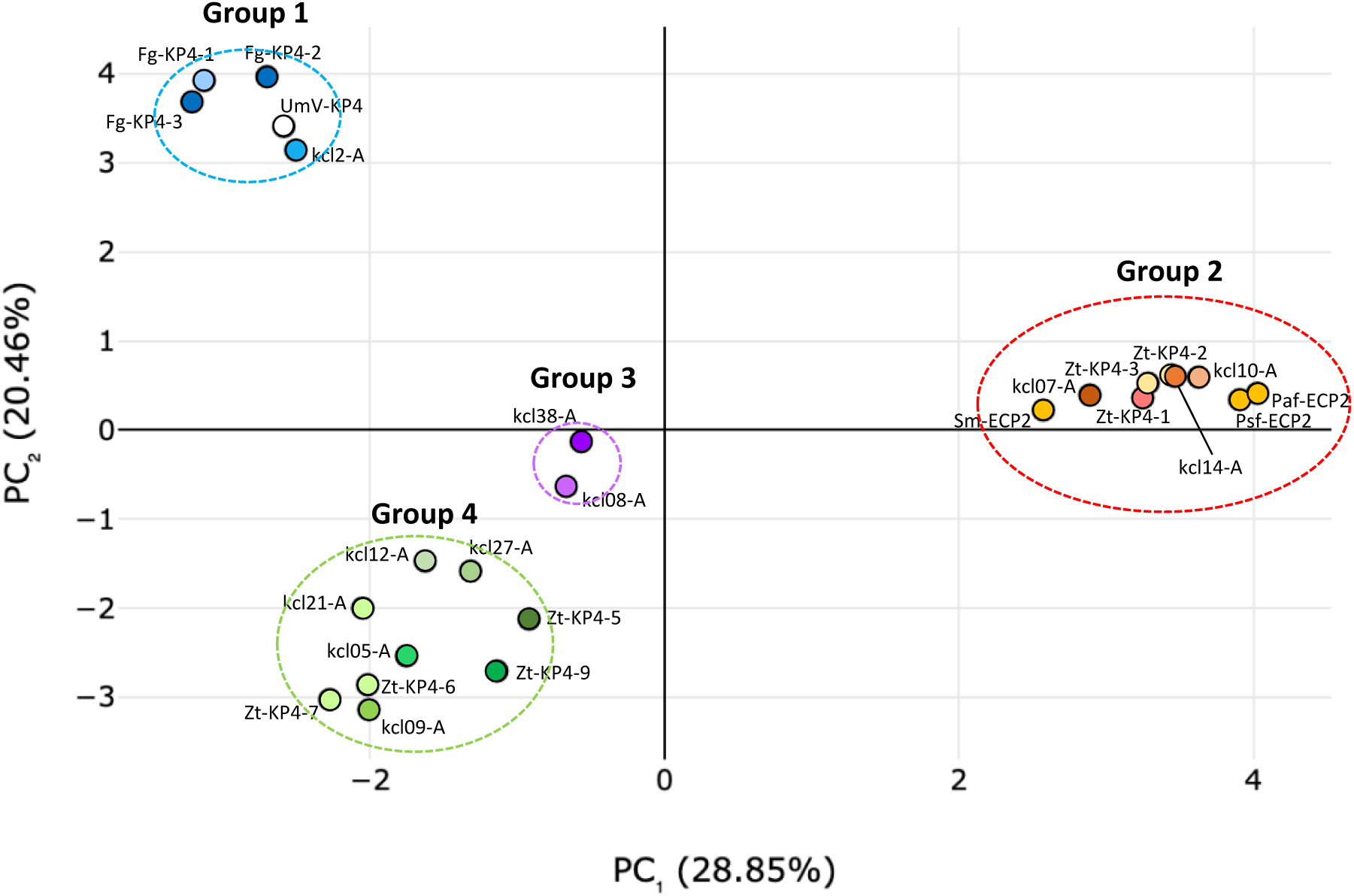
Classification of fungal KP4 families according to their structures. DALI all by all analysis followed by a PCA using a DALI score matrix [28]. meta-cluster kclXX, AFDB50 cluster kclXX-Y, (number of proteins in the AFDB50 cluster), Uniprot number of the representative. kcl01-D, (83), Zt-KP4-1 = A0A2H1H404/8ACX; kcl01-Q, (203), Sm-ECP2 = N1QK49, Psf-ECP2 = M2Z6V3, Paf-ECP2 = Q00365; kcl02-A, (15), A0A4U0XEI9; kcl02-E, (256), Fg-KP4-2 = A0A3S7E9S6, Fg-KP4-3 = A0A2H3GQS6; kcl02-I, (125), Fg-KP4-1 = A0A2H3GK41; klc02-0, (324), Zt-KP4-9 = A0A2H1GXX1; kcl05-A, (14), A0A6A6DAE0; kcl07-A, (53), A0A177DVW3; kcl07-C, (35), Zt-KP4-2 = A0A2H1FWY7; Zt-KP4-3 = A0A1Y6M296; kcl08-A, (5), S3D5P; kcl09-A, (14), M3BT27; kcl09-B, (131), Zt-KP4-6 = A0A2H1GNH3, Zt-KP4-7 = A0A2H1GND9; kcl10-A, (16), A0A1L9SH47; kcl12-A, (9), E3L0Y8; kcl14-C, (12), A0A0A1V5R3; kcl18-A, (9), Zt-KP4-5 = A0A2H1GP20; kcl21-A, (13), A0A0J9W663; kcl27-A, (39), A0A4T0WCM2; kcl38-A, (8), A0A017SS08. White circle, UmV-KP4 = kptA

## DISCUSSION

The recent systematic investigation of the structures of fungal effectors revealed extended protein families with conserved structures [5,20,21]. In this study, the elucidation of the structure of two *Z. tritici* effectors (Zt-Mycgr3-91409-2, Zt-NIP1) by X-ray crystallography revealed their unexpected structural similarity to the UmV-KP6α and UmV-KP4 killer toxins from the dsRNA virus infecting the fungal pathogen *U. maydis* [26]. Both effectors, renamed Zt-KP4-1 and Zt-KP6-1, were toxic to fungi, but not to wheat. We established that proteins structurally related to Zt-KP4-1 and Zt-KP6-1 are widespread and abundant in fungi, and belong to extended super families of structurally related proteins. These results provided a novel comprehensive classification of these effectors into, respectively, four KP4 and three KP6 super families, providing a solid framework for studying these fascinating proteins.

### *UmV*-KP6 and *UmV*-KP4 killer toxins

*U. maydis* secretes three killer proteins (UmV-KP1, KP4 and KP6) encoded by double-stranded RNA viruses (UmV) [26,47,48]. They kill susceptible *U. maydis* strains, while UmV-infected strains are resistant to the killer protein they produce [25,26]. UmV-KP4 inhibited fungal growth presumably by perturbing cellular calcium homeostasis [48], and inhibited L-type voltage-gated (LVG) Ca^2+^ channels in mammalian cells [49]. Interestingly, UmV-KP4 also perturbed Ca^2+^ gradients of plant root hair cells [50]. The *P. patrens* moss KP4 protein also modified Ca^2+^ gradients of moss cells [51]. However, UmV-KP4 does not induce major growth defects, necrosis, chlorosis or cell death when over-expressed in transgenic plants such as wheat, corn or tobacco [52–55]. UmV-KP6α and UmV-KP6β associate to form a stable heterodimer [29]. UmV-KP6α appears necessary for recognition of susceptible cells, through a mechanism involving the N-terminal α helix, while UmV-KP6β is required for toxicity [26]. However, the molecular mechanism underpinning the toxicity of UmV-KP6α/β to fungal cells is unknown [26], although it has been hypothesised that this complex forms membrane pores [29]. These previous studies on KP4 and KP6 proteins provided interesting insights into their function. However, a full understanding of their mode of action on plant and fungal cells and their biological function is still lacking.

### Zt-KP6-1 and Zt-KP4-1 are antifungal proteins

Neither Zt-KP4-1 nor Zt-KP6-1 induced necrosis when infiltrated into wheat leaves. Similarly, Fg-KP4L-2 from *F. graminearum* did not induce necrosis of wheat leaves [46]. Still, recombinant Fg-KP4L-2 slightly inhibited the growth of wheat roots [46]. This is reminiscent of the low activity of UmV-KP4 on *A. thaliana* root growth [50]. We showed that both Zt-KP4-1 and Zt-KP6-1 inhibited the growth of *Z. tritici* and *B. cinerea*. Within *U. maydis*, UmV-KP4 and UmV-KP6 were only toxic to susceptible strains [26]. However, UmV-KP4 inhibited the growth of other fungi such as *Tilletia caries*, *Ustilago tritici*, *Alternaria alternata* and *Phoma exigua* [54], but not the yeast *S. cerevisae* [56]. When overexpressed in transgenic wheat or tobacco, UmV-KP4 conferred resistance to, respectively, *T. caries* and *U. tritici* [52,53,55], and *A. alternata* and *P. exigua* [54]. UmV-KP6 was also toxic to the yeast *Brettanomyces bruxellensis* [57], but not *S. cerevisae* [47,57]. The structure-activity relationship of Zt-KP6-1 likely differs from UmV-KP6α/β as it did not require the formation of a heterodimer to be toxic [26,29]. In addition, Zt-KP6-1 does not have the N-terminal α-helical extension of UmV-KP6α critical for its toxicity [26,29]. Since structural homologs of Zt-KP4-1 and Zt-KP6-1 are widespread in fungi, it is tempting to speculate that other KP4 and KP6 proteins are also toxic to fungi. Recent work on plant pathogenic fungi highlighted effectors with antimicrobial activities [58]. For instance, the broad host range plant pathogenic fungus *Verticiulium dahliae* secretes several effectors during host xylem invasion that have an antibacterial activity [59]. These antimicrobial effectors changed the composition of the xylem and root bacterial microbiota of host plants [60,61]. In particular, VdAve1, an effector with antibacterial activity, specifically inhibits the growth of *Sphingomonas* bacteria, which acts antagonistically toward *V. dahliae* in the host plant rhizosphere [59,62]. AMAPEC, a machine-learning based computational software using shared structural and physicochemical properties of known antimicrobial proteins, identified a large number of candidate antimicrobial proteins among effectors of three different plant-associated fungi [63]. This finding suggests that fungi have extended repertoires of antimicrobial effectors. Most of KP4 and KP6 proteins identified in this study were predicted to be antimicrobial according to AMAPEC (83% of KP6, 77% of KP4, Sup Table 5), including Zt-KP4-1 and Zt-KP6-1. It will therefore be interesting to investigate whether KP4 and KP6 super families are primarily composed of antimicrobial effectors.

### Fungal KP4 and KP6 proteins are widespread in fungi and define novel structural super families with possible conserved functions

To identify structural homologs of Zt-KP4-1 and Zt-KP6-1, we developed a novel pipeline based on (i) FoldSeek structure similarity searches using AF2 structural models of fungal secreted proteins, combined with (ii) HMM pattern searches relying on Zt-KP4/UmV-KP4 or Zt-KP6/UmV-KP6α conserved cysteine patterns and (iii) structure-based manual curation (FHAT, Sup Figure 9). In this last important step, the topology and structure of KP4 and KP6 proteins were manually compared to Zt-KP4/UmV-KP4 or Zt-KP6/UmV-KP6α. This procedure discriminated proteins with a true KP4 or KP6 structural fold from proteins with limited or no structural similarity. To assess the performance of FHAT, we compared our dataset with those of two recent systematic structural studies of fungal effectors [20,21]. Clustering the KP4 and KP6 proteins identified in the three studies into families of highly related proteins (at least 90% sequence identity and 60% coverage) identified 1737 KP4 families for FHAT, 227 for Seong and 333 for Derbyshire, as well as 559 KP6 families for FHAT, 97 for Seong and 80 for Derbyshire. The differences in the numbers of proteins identified mostly come from large differences in the number of fungal genomes analyzed. We therefore specifically compared these datasets for proteins detected in the seven shared fungal species (Figure 7). Most KP4 and KP6 identified by FHAT in *Z. tritici* were also found by Seong and/or Derbyshire (80%, Figure 7). However, FHAT was the only method uncovering Zt-KP4-1, Zt-KP4-5 and Zt-KP6-6. Ten KP4 and two KP6 were specifically detected by Derbyshire in *Z. tritici* (Figure 7). However, their structures did not display the typical KP4 or KP6 fold, suggesting that these proteins were false positive. No false positive were detected by Seong in *Z. tritici*. The same trend was observed for the six other shared species (Figure 7). This result suggested that FHAT was more sensitive than previous analyses, and has a low false positive rate. Consequently, it provides a comprehensive inventory for these important effector families in fungi.

**Figure 7.**
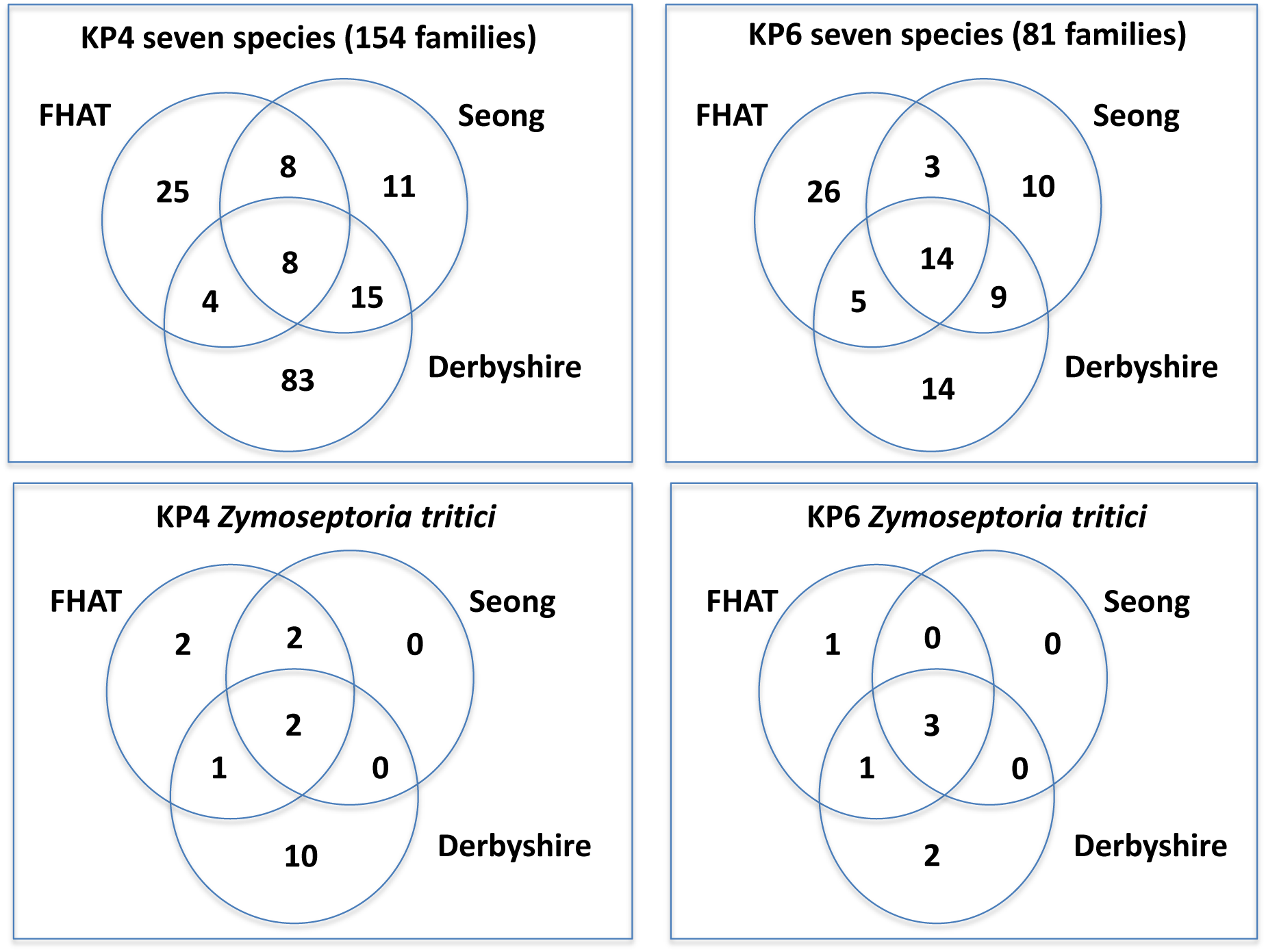
KP4 and KP6 proteins identified by FHAT and previous methods. FHAT (Sup. Figure 10), Seong: [20], Derbyshire: [21]. KP4 and KP6 protein sequences found in the seven shared fungal species by the three methods, were clustered into highly related protein families sharing at least 90% sequence identity and 60% sequence overlap. The numbers of effector families detected by one or more methods are indicated in the Venn diagrams for KP4 and KP6 proteins of either the seven shared species or only *Z. tritici*. The seven shared species are: *Blumeria graminis*, *Botryotinia fuckeliana, Fusarium oxysporum, Leptosphaeria maculans, Magnaporthe oryzae, Verticillium dahliae, Zymoseptoria tritici*.

KP4 and KP6 proteins were found in both *ascomycotina* and *basidiomycotina* with different lifestyles, including plant pathogens, plant symbionts and saprophytes, but excluding as a notable exception animal fungal pathogen. Contrary to KP6 proteins, KP4 proteins were identified outside fungi, in plants. Some plant KP4 clustered with Fg-KP4 and UmV-KP4 in AFDB cluster kcl02-E from Group-1 (Figure 6), as observed by Brown (2011), Lu and Faris (2019), and Guan (2023). Another set of plant KP4 (175 proteins) were related at the sequence level since they belonged to the same AFDB cluster (kcl01-R) and they shared the same pfam domain (PF1657) characteristic of plant antifungal proteins such as Ginkgobilobin-2.

Our structure-based classification of the KP4 structural family integrated results from previous large-scale comparative studies and created a robust framework for future research (see Figure 6). KP4 structural Group-1 overlapped with the KP4 family characterized by the presence of a KP4 pfam domain defined using Fg-KP4 and UmV-KP4 by Lu and Faris (2019), and Brown (2011). KP4 structural Group1 contained 396 KP4 proteins from AFDB clusters kcl02-A, kcl02-E and klc02-I including 3 KP4 of the moss *Physcomitrium patens* from kcl02-E [51]. KP4 structural Group-2 overlapped with the Ecp2 family characterized by the presence of a HCE2 pfam domain [39]. KP4 structural Group-2 contained 402 proteins from the clusters kcl01-D, kcl01-Q, kcl07-A, kcl10-A and kcl14-A. The discovery that KP4 and Ecp2 families are major groups of the KP4 structural family is a novel finding generating far-reaching hypothesis on their function and evolution. Combined with the finding that UmV-KP4 and Zt-KP4-1 from structural Group-1 and -2 are toxic to fungi, we hypothesized that KP4 proteins from these two structural groups are antimicrobial. Such effectors could be essential for the competition of fungi during their colonization of particular ecological niches. This hypothesis could be extended to Group-3 and -4 KP4 even though they neither contained HCE2 or KP4 domains, nor have known biological function. The following step will be to investigate the antimicrobial activity of a wide array of different KP4 proteins.

Our structure-based classification of the KP6 structural family provided a unifying framework facilitating future research on these proteins (Figure 5). The KP6 structural Group-1 included AvrLm10A and SIX5 that physically interact with effectors encoded by neighbouring genes, facilitating their symplastic transport in host plant tissues [14,16,64,65]. We hypothesized that his group, containing 25 fungal proteins from the kcl04-C AFDB cluster but no proteins from *Zymoseptoria*, has evolved to act in the cell-to-cell transport of other effectors [14]. The situation was different with proteins from KP6 structural Group-2 and 3. KP6 structural Group-2 included the antifungal protein UmV-KP6β, the known effectors BAS4 from *P. oryzae* [43] and ECP28 from *P. fulvum* [42], Zt-KP6-4, and 420 KP6 proteins from clusters kcl01-C, kcl02-G, kcl04-A and kcl11. Similarly, KP6 structural Group-3 contained the two antifungal proteins Zt-KP6-1 and UmV-KP6α, as well as Zt-KP6-2, Zt-KP6-6, and five KP6 proteins from cluster kcl02-A. We hypothesized that these two KP6 structural Groups could gather proteins with antimicrobial activity. Testing this hypothesis will require the investigation of the antimicrobial activity of a wide array of KP6 proteins.

## MATERIELS AND METHODS

### Protein expression and purification

The Zt-KP6 mature protein (residues 21-90) was expressed with the pET-SP vector, produced and purified as previously described [5]. The Zt-KP4 protein was expressed with the pET derived plasmid pdbccdb-3C-His, produced and purified as previously described [7] (Supplementals File S1).

### Nuclear magnetic resonance spectroscopy

To generate isotopically labelled samples for NMR spectroscopy, we used ^15^NH_4_Cl, as the unique source of nitrogen. The samples containing 0,7 mM of purified protein in 10 % D_2_O and 1 μM DSS as a reference were analyzed using an 800 MHz Avance Bruker spectrometer and a TCI probe at 303 K. Spectra were recorded with Topspin 3.2 (Bruker) and referenced to DSS (4,4-dimethyl-4-silapentane-1-sulfonic acid) for the ^1^H dimension and indirectly for the ^15^N dimension. NMR data were processed using Topspin 3.6 (Bruker) and analyzed with the in-house software Cindy 2.1 (Padilla, www.cbs.cnrs.fr). Side chain assignments were carried out using 2D-NOESY, 2D-TOCSY and COSY-DQF experiments with D_2_O samples, combined with ^15^N-NOESY-HSQC and ^15^N-TOCSY-HSQC 3D spectra. The CYANA 3 program [66] was used for automatic NOE assignments and structure calculations. The CANDID procedure of CYANA was used to assign the 3D-peaks list from the ^15^N-NOESY-HSQC spectra. Structures were analyzed with PROCHECK [67]. Refinement statistics for the structure are listed in Sup. Table 1. The structure of Zt-KP6 was deposited in the Protein Data Bank and in BMRB with accession number 9GWD and ID 34963 respectively.

### X-ray crystallography

The purified protein Zt-KP6 was concentrated to 10 mg/ml in 20 mM Tris-HCl, pH 8.0, 150 mM NaCl, 1 mM DTT. The purified protein Zt-KP4 was concentrated to 8 mg/ml in 20 mM NaCi, pH 5.6, 150 mM NaCl, 1 mM DTT. Crystallization trials were performed at 20°C by the hanging-drop vapor-diffusion method in 96-well microplates using a Mosquito HTS robot with 100 nl of protein mixed with 100 nl of reservoir. After 3 weeks, Zt-KP6 crystals were obtained from the condition 0.01 M NiCl_2_, 0.1 M Tris pH 8.5, 1M LiSO_4_ (from classic kit, Hampton research®). After refinement and soaking in tantalum, diffraction crystals were obtained in 0.1M Tris pH 9, 1.5M LiSO_4_, 100mM ZnSO_4_. After 2 weeks, small crystals of Zt-KP4 were obtained from one condition of the Peg kit (Qiagen®) containing 0.1M sodium acetate pH 4.6 and 30%(v/v) PEG 300. All crystals were flash frozen in liquid nitrogen.

### Data collection, processing and structure determination

Crystal diffraction datasets were collected at the European Synchrotron Radiation Facility (ESRF, Grenoble). For Zt-KP6, data were collected at beam line ID30A-1 [68–72] using a Pixel detector (Pilatus3-2M) and processed by XDS [73] and AIMLESS [74], from the CCP4/CCP4i programs suite [75,76]. For Zt-KP4, data were collected at beam line ID30B [77,78] using a Pixel detector (PILATUS3 6M) and auto processed by XDSAPP [79,80]. Zt-KP6 crystals belong to the *P*2_1_2_1_2_1_ space group and contain two molecules in the asymmetric unit. The structure of Zt-KP6 was determined to 1.36 Å by the single wavelength anomalous diffraction (SAD) method using HKL2MAP [81,82]. Zt-KP4 crystals belong to the H3 space group and contain eight molecules in the asymmetric unit. The structure of Zt-KP4 was determined by molecular replacement with Molrep [83] from CCP4I2 [76] using the model determined by AF2 [19]. After modelbuilding using arp/warp [84], Coot [85] and refinement by REFINE from PHENIX [86,87] final structures had R(%) / R(%)free of 14 / 18.3 for Zt-KP6 and 17.5 / 21.9 for Zt-KP4. Data collection and refinement statistics for the structures are listed in Sup. Table 2. Figures were generated using PyMol v1.6 (http://www.pymol.org/). The atomic coordinates and structure factors of Zt-KP6 and Zt-KP4, were deposited in the Protein Data Bank with accession numbers 6QPK and 8ACX respectively.

### Biological assays of Zt-KP4-1 and Zt-KP6-1

Zt-KP4-1 and Zt-KP6-1 were infiltrated into leaves of the wheat cultivar Taichung 29 at a concentration of 2 µM. As a control, the necrotrophic effector ToxA from *P. nodorum* [33] was infiltrated into Taichung 29 wheat leaves at concentrations from 0.2 to 2 µM. *Z. tritici* IPO323 isolate and the necrotrophic plant pathogen *Botrytis cinerea* were used for growth assays in 96 well microtiter plates containing liquid potato dextrose growth medium (PDA). Each well of the plate was filled with 100µL of PDA containing either 10E6 *Z. tritici* spores/mL or 10E6 *B. cinerea* spores/mL and different concentrations of Zt-KP4-1 or Zt-KP6-1 (0.55, 1.65 and 5.5 µM). 96 well plates were incubated in the dark at either 18°C (*Z. tritici*) or 25°C (*B. cinerea*) with shaking (50 rpm). After 7 days of growth, the OD at 580 nm was recorded on a spectrophotometer. The % of inhibition was calculated according to the formula: % inhibition = ((OD treated/OD control)-1)*100. Values displayed in Table 1 were the mean ± Standard Error Deviation (SED) from three independent experiments.

### FHAT protocol for the detection of fungal KP4 and KP6 proteins

Fungal KP4 and KP6 proteins were retrieved from public databases according to the following protocol (Sup Figure 9) and result files for each step are available at URLs https://pat.cbs.cnrs.fr/kp4/fhat_kp4 and https://pat.cbs.cnrs.fr/kp6/fhat_kp6. The protocol for the detection of fungal KP4 and KP6 proteins is fully described in the legend of Sup Figure 9.

### PCA analysis

To classify KP4 and KP6 proteins according to their structural similarities, an “all against all” analysis was performed for both families using the DALI server [28]. This computing delivered matrices of all the Z-scores for each pair of proteins. If a pair of proteins had a Z-score of 0.1, they were excluded from the following step. A Principal Component Analysis (PCA) was performed using these Z-scores matrices using *Statistic Kingdom* (https://www.statskingdom.com/410multi_linear_regression.html). Protein names were directly labeled on the graphs.

## Acknowledgements

The CBS is a member of the French Infrastructure for Integrated Structural Biology (FRISBI), national infrastructure supported by the French National Research Agency (ANR-10-INBS-0005). We thank R. Lambert for assistance with protein production and x-ray crystallography. We are grateful to Matthew Bowler and Didier Nurizzo at the European Synchrotron Radiation Facility (ESRF), Grenoble, France for providing assistance in using beamline ID30B and ID30A-1 and the ESRF for provision of synchrotron radiation facilities via Block Allocation Group beamtime.

## Data availability

The data that support the findings of this study are openly available at the following URLs:

https://pat.cbs.cnrs.fr/kp4/fhat_kp4

https://pat.cbs.cnrs.fr/kp6/fhat_kp6

## Authors contributions

KDG, AP, TK and MHL planned and designed the research. KDG, LM, JG, PB, FH, ML, JR, YPH and MHL performed experiments and analysed data. KDG, AP, TK and MHL wrote the manuscript.

## List of supplementary figures and tables

**Supplementary Figure 1. Annotation of IPO323 *Z. tritici* Zt-Mycgr3-91409 (Zt-KP6-1) and Zt-Mycgr3-106176 (Zt-KP4-1)**

Sup. Fig.1-A Annotation of IPO323 *Z. tritici* CDS Zt-Mycgr3-91409 (Zt-KP6-1)

Sup. Fig.1-B Annotation of IPO323 *Z. tritici* CDS Zt-Mycgr3-106176 (Zt-KP4-1)

**Supplementary Figure 2 Purification of Zt-KP6-1 by gel filtration**

Gel filtration profile with a Superdex S75 26/60 (GE Healthcare) column at 2.5 ml/min in buffer (20 mM Tris-HCl, pH 8.0, 150 mM NaCl, 1 mM DTT) and SDS-PAGE analysis of purified Zt-KP6-1. Sizes of the molecular mass markers are indicated on the left of the gel.

**Supplementary Figure 3 NMR analysis of Zt-KP6-1**

HSQC spectra of Zt-KP6-1 at, 0.7 mM in 20 mM NaCitrate pH 5.4, 150 mM NaCl, 1 mM DTT, 303 K, 800 MHz. Cross-peak assignments are indicated using one-letter amino acid and number (the asterisk shows a folded red peak). Missing residue N36.

**Supplementary Figure 4 Topology diagrams of Zt-KP6-1 and UmV-KP6α**

The structures were processed by PDBsum and the secondary structures were colored from N-terminus (blue) to C-terminus (red) according to Figure 1. Disulphide Bonds (D) are represented as grey lines. The disulphide pattern of the UmV-KP6 proteins starts with a CxC cysteine motif in β1 bound to cysteine residues in, respectively, β4 and H2 (equivalent to H1 in ZtKP6-1). The first cysteine in the β1 CxC motif and the linked cysteine in β4 were missing in Zt-KP6-1 without affecting the antiparallel arrangement of β4 and β1.

**Supplementary Figure 5 Purification of Zt-KP4-1 by gel filtration**

Gel filtration profile with a Superdex S75 16/60 (GE Healthcare) column at 1 ml/min in buffer (20 mM NaCi, pH 5.6, 150 mM NaCl, 1 mM DTT) and SDS-PAGE analysis of purified Zt-KP4-1. (a) dimer, (b) monomer. Sizes of the molecular mass markers are indicated on the left of the gel.

**Supplementary Figure 6 Dimer of Zt-KP4-1 between chains B (cyan) and G (blue)**

Residues involved in hydrogen bonds at the dimer interface are labelled and represented by sticks. Hydrophobic residues involved in the pocket are highlighted in lines and coloured in yellow. According to the GenBank sequence SMR60557.1, the residue numbering must start after the signal peptide at residue S20. T33 in the structure corresponds to T48 in the GenBank sequence. Binding is mediated by four hydrogen bonds between the Thr48, Ser50, Gln90, and Thr158 from chain E, G, F or C and Thr158, Gln90, Ser50 and Thr48 from chain H, B, C or A respectively, as well as a hydrophobic pocket formed by residues Val62 and Val149 of both chains.

**Supplementary Figure 7 Topology diagrams of Zt-KP4 and UmV-Kp4**

The structures were processed by PDBsum and the secondary structures were coloured from N-terminus (blue) to C-terminus (red) according to the Figure 2. Disulphide Bonds (D) are represented as grey lines. For UmV-KP4, the five disulphide bonds were located between Cys5 in the N-terminus and Cys78 at the end of helix H3, between Cys11 in H1 and Cys81 just before the start of β5, between Cys27 in H2 and Cys67 in H3, between Cys35 at the end of β1 and Cys60 at the start of β4 and finally between Cys44 at the end of β2 and Cys105 at the C-terminus (Figure 2). In their study, Gage et al. (2002) reported that changing Lys42 of UmV-KP4 to glutamine reduced KP4 biological activity by 90%. In Umv-KP4, Lys42 stabilised the disulphide bridge involving Cys44 and Cys105 and it was proposed to be required for the stability of the C-terminus (Gage et al. 2002). In Zt-NIP1, there was no amino acid involved in stabilizing the C-terminus and the amino acid corresponding to Lys42 was replaced by Thr85. However, the C-terminus of Zt-NIP1 was stabilised by the β7 strand.

**Supplementary Figure 8 Structure and topology of Zt-KP6-1 and Zt-KP6-2**

Zt-KP6-2 structure was predicted by AlphaFold2. The structures were processed by PDBsum and the secondary structure were coloured from N-terminus (blue) to C-terminus (red) according to Figure 1.

**Supplementary Figure 9 Flow chart of the pipeline used to identify fungal proteins structurally related to Zt-KP6-1 and Zt-KP4-1**

- Step 1: Selection of the reference templates (result file1): First, PDB structures were selected as reference templates for the validation of candidate 3D folds (1KPT-A and 8ACX-A in the case of KP4, 4GVB-A and 6QPK-A in the case of KP6).

- Step 2-1: Foldseek search (result files 2 and 4): AlphaFold2 models similar to the reference PDB templates were detected in the AFDB50 database using the Foldseek server (van Kempen et al. 2024).

- Step 2-2: The fungal representatives of the AFDBcluster database (Barrio-Hernandez et al. 2023) were collected (result file 6) if they shared a Foldseek E-value below 1e-3 with any hit from the previous Foldseek search.

Output 1: AFDB50 cluster representatives corresponding to the KP4 and KP6 candidate proteins detected by Foldseek and AFDBcluster database.

- Step 3: Hmmer search (result file 3 and 5): The KP4 and KP6 proteins detected by Foldseek in step 2-1, whose TM scores were above 0.5 were aligned according to TMalign output. The resulting multiple sequence alignment was used as input query to search for homologous sequences shorter than 300 residues using 10 Hmmer iterations with E-value<1 and query overlap >75% in a custom fungal secretome database. This custom database was composed of 713,879 secreted fungal proteins which were predicted using SignalP 5.0 (Almagro Armenteros et al. 2019) from 9,785,231 protein sequences originating from 1014 annotated fungal genomes available at the Ensembl web site (ftp.ensemblgenomes.org release-46).

Output 2: AFDB50 cluster representatives corresponding to the KP4 and KP6 candidate proteins detected by Foldseek and Hmmer.

- Step 4-1: Filtering (result file 8): The representatives of the AFDB50 clusters covering all candidate proteins detected at both Step 2 (Output-1, Foldseek) and Step 3 (Output-2, Hmmer) were selected if they were not predicted as transmembrane and if their sequences were shorter than 180 residues. The members not predicted as secreted by SignalP 5.0 were removed (Almagro Armenteros et al. 2019).

- Step 4-2: Structural validation (result files 9 to 12): The AlphaFold2 models of all the representatives of the selected AFDB50 clusters, were validated if their TM-score was above 0.5 when superposed by TMalign onto one the reference templates (Zhang et Skolnick 2005). The TMscore cutoff was set at a rather high value of 0.5 to minimize the detection of false positives.

Output-3: Members and representatives of the KP4 and KP6 AFDB50 clusters (kclX-Y) filtered using Step4 criteria. Members = result file 18, Representatives = result file 13

- Step 5: Structural alignment (result file 15 and 16): Representatives of KP4 and KP6 AFDBclusters were structurally aligned using TMalign onto a reference PDB template, respectively on chain A of the PDB structure 1KPT for KP4, and on chain A of the PDB structure 4GVB for KP6. Protein secondary structures were added at the end of each alignment. Same as Step 4.3: Phylogeny (result file 17): Phylogenetic trees of Representatives of KP4 and KP6 AFDBclusters were constructed using FastME (Lefort, Desper, et Gascuel 2015) using structural alignments after removing the aligned positions with more than 50% indels. Taxonomic information was suffixed to each protein identifier.

- Step 6: Uniprot annotations analysis (result file 19 and 20): The most frequent Uniprot annotations (UniProt Consortium, 2023) and the taxonomic distributions of the two validated member sets were computed for further analysis. Annotations were listed in a web page hyper-linked to external databases for an easy analysis of the proteins whose domains, structures, orthologous groups or functions are already known. A subset analysis was performed (result file 34 to 38): The most frequent Uniprot / Pfam annotations of the protein subsets were collected. The AlphaFold2 models of each subset protein was aligned using TMalign onto 8ACX for the KP4 like subset and 6QPK for the KP6 like proteins. FastME trees were derived from these structural alignments after the removal of their aligned positions with more than 50% indels. Known Pfam family identifiers were suffixed to each protein identifier in the last result tree. Lastly, each pair of selected 3D models was superposed using TMalign and a structural tree was inferred from the obtained TM-scores using FastME. This tree was compared with a phylogenetic tree derived from the corresponding multiple sequence alignment built using MAFFT.

- Step 7: *Zymoseptoria* analysis (result file 22 to 24): The KP4 like and KP6 like detected in *Zymoseptoria* were superposed onto experimental structures 8ACX and 6QKP. The TMscores of each superposition were collected and phylogenetic trees were inferred using TMalign from both *Zymoseptoria* alignments without the columns with more than 50% indels.

- Step 8: Comparison of the final set to previous studies (result file 25 and 26): The detected KP4 like set was compared to those detected in previous studies on fungal effectors (Seong et Krasileva 2023; Derbyshire et Raffaele 2023). By using Kclust (Hauser, Mayer, et Söding 2013), the validated KP4 like proteins were clustered with the KP4 proteins detected by the other studies if their sequences shared more than 30% sequence identity and more than 80% sequence overlap. The diversity of the KP4 like protein set detected by each method was estimated as the percentage of the above 30%id-max clusters which include at least one protein. The same clustering analysis was done for the KP6 proteins.

- Step 9: K-clust Clustering (result file 29 to 33): We have clustered all the proteins from Step 4 (Representatives of the selected KP4 and KP6 AFDBclusters) using Kclust at 40% maximum shared sequence identity. Meta-clusters (kclX) were selected by a hierarchical clustering of the representatives which share more than 40% identical positions between both their amino acid sequences and their secondary structures coded as 3-classe sequences (H for helix, E for extended strand, C for coil).

- Step 10: Final structural validation. The structures of the proteins representative of each meta-cluster from Step 5 (kcl-X) were predicted by AlphaFold2. If the representative of the meta-cluster displayed a pLDDT score below 50, another member of the same meta-cluster with pLDDT above this value was used.

The topology of each KP4 representative was visually inspected with Pymol, and it was validated or rejected as a KP4 according to following topology: H2-β1-β2-β3-β4-H3-β5-β6. The first 15 N-terminal residues were not taken into account. To be validated the candidate protein must display two antiparallel alpha-helices H2 and H3. The two antiparallel strands β1 and β4 were sometime poorly defined, since these two strands were short and their arrangement difficult to confirm. Strands β2 and β3 must be present. However, β2 was sometime replaced by an alpha-helix. The presence of the antiparallel strands β5 and β6 as well as β6-β3 and their proximity must be present. Candidate KP4 proteins that do not meet these criteria were eliminated from further consideration.

The topology of each KP6 representative was visually inspected with Pymol, and it was validated or rejected as a fold according to following topology: H2-β1-β2-β3-β4-H3-β5-β6. The validation of a KP6 fold required the following topology: β1-H2-β2-β3-H3-β4. The KP6 fold required for validation was defined according to the following topology: β1-H2-β2-β3-H3-β4. The short N-terminal H1 helix does not play a role in determining the core topology, and it was not taken into account. The two antiparallel helices H2 and H3 were detected in the structure of all the selected KP6 meta-cluster representatives. To be validated the candidate protein must display antiparallel strands β1, β3 and β4 with β1 strand central and in antiparallel arrangement with β3 and β4. The two antiparallel strands β2 and β3 must be present and H3 must be connected to β4. Overall, the β sheet formed by β1-β2-β3-β4 must be observed on one side (sometimes with poorly defined strands) and the two helices H2 and H3 on the other side. Candidate KP6 proteins that do not meet these criteria were eliminated from further consideration.

- Step 11: Final members and representatives of KP4 or KP6 clusters

The result file(s) of each analysis step are displayed for both KP4 and KP6 protein families at the following URLs:

- https://pat.cbs.cnrs.fr/kp4/fhat_kp4

- https://pat.cbs.cnrs.fr/kp6/fhat_kp6

**Supplementary Figure 10 Structure and topology of Zt-KP4-1, Zt-KP4-2 and Zt-KP4-3**

Zt-KP4-2 and Zt-KP4-3 structures were predicted by AlphaFold2. The structures were processed by PDBsum and the secondary structures were coloured from N-terminus (blue) to C-terminus (red) according to Figure 2.

**Supplementary Figure 11 Expression of Zt-KP4-1, Zt-KP4-2, Zt-KP4-3, Zt-KP6-1, Zt-KP6-2, Zt-KP6-4 and Zt-KP6-6 during infection of wheat leaves infection.**

RNAseq data from Rudd et al. (2015). RPKM (fungal RNAseq Reads Per Kilobase Million)

**Supplementary Table 1. X-ray data collection and refinement statistics for Zt-KP6-1 and Zt-KP4-1**

**Supplementary Table 2. NMR and refinement statistics for Zt-KP6-1**

(Zt-KP6-1, 0.7 mM, 20 mM NaCitrate pH 5.4, 150 mM NaCl, 1 mM DTT, 303 K, 800 MHz) PDB:9GWD BMRB ID 34963

**Supplementary Table 3 List of the proteins structurally related to Zt-KP6-1 detected using our pipeline**

**Supplementary Table 4 List of the proteins structurally related to Zt-KP4-1 detected using our pipeline**

**Supplementary Table 5 Prediction of the antimicrobial activity of fungal KP4 and KP6 proteins using AMAPEC** [88].

**Supplementary file S1 Nucleotide sequence of synthetic genes and plasmids used in this study**

## Notes

https://pat.cbs.cnrs.fr/kp4/fhat_kp4

https://pat.cbs.cnrs.fr/kp6/fhat_kp6

### Competing Interest Statement

The authors have declared no competing interest.

### Summary of Updates

Title review; author affiliations updated; Updated the section on the distribution of KP4 and KP6 in fungi for clarity and the discussion section.

