## supplementary figures table for "*Zymoseptoria tritici* effectors structurally related to killer proteins UmV-KP4 and UmV-KP6 are toxic to fungi, and define extended protein families in fungi"

**Supplementary Figure 1. Annotation of IPO323 *Z. tritici* Zt-Mycgr3-91409 (Zt-KP6-1) and Zt-Mycgr3-106176 (Zt-KP4-1)**

**Sup. Fig.1-A Annotation of IPO323 *Z. tritici* CDS Zt-Mycgr3-91409 (Zt-KP6-1)**

Sup. Fig.1-A1 Annotation of Zt-Mycgr3-91409 (Zt-KP6-1, chr_3:487000-490000) using transcript evidence (Lapalu et al., 2023)

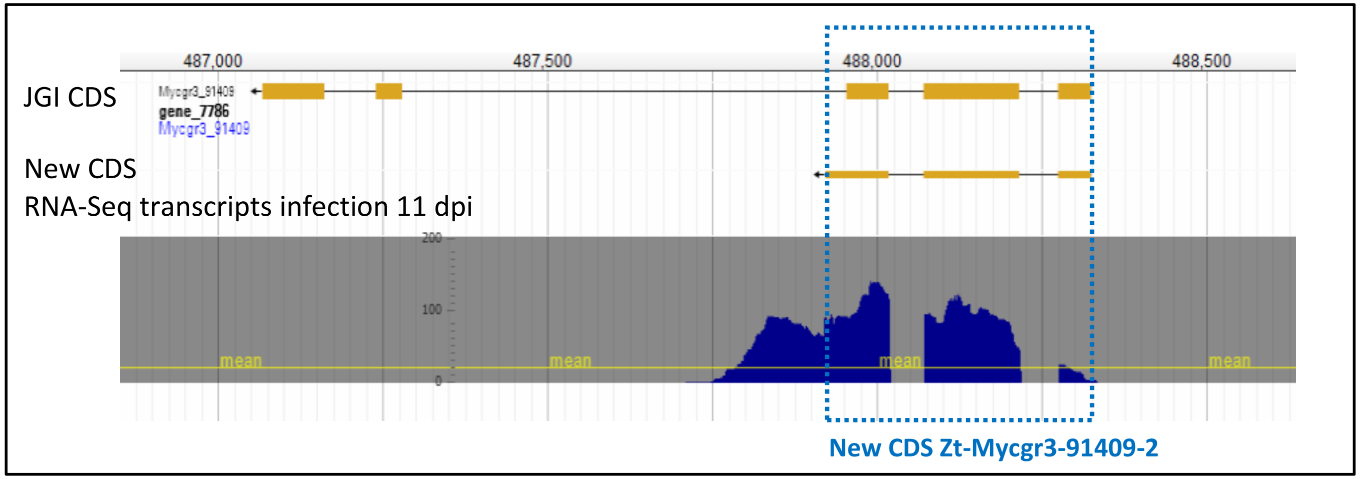


Sup. Fig.1-A2 Protein sequences of Zt-Mycgr3-91409 and Zt-Mycgr3-91409-2

>Zt-Mycgr3-91409
MAPIFTYAVAALAFAQSAYAVVYAARCKFGNPLVQNNRITRAVCDLTNEHTTKDGSWHYVEVDNECKYLAGDNPRDQPGWAVFVK
YCPEQNAAADKSRARGEQTCGFVRTPVDDVSRRAESATTPVCEGD

>Zt-Mycgr3-91409-2(Zt-KP6-1)
MAPIFTYAVAALAFAQSAYAVVYAARCKFGNPLVQNNRITRAVCDLTNEHTTKDGSWHYVEVDNECKYLAGDNPRDQPGWAVFVK
YCTYYKGVPDA

Sequences identical in Zt-Mycgr3-91409 and Zt-Mycgr3-91409-2 are highlighted in blue

Sup. Fig.1-A3 Proteins similar to Zt-KP6-1 in annotated *Z. tritici* genomes.

SMY22152.1 (Uniprot A0A1Y6LCD6), *Z. tritici* strain 1A5
SMQ48456.1 (Uniprot A0A1X7RM13), *Z. tritici* strain 3D7
SMR48303.1 (Uniprot A0A2H1G421), *Z. tritici* strain 1E4, PDB_6QPK
SMR49455.1 (Uniprot A0A2H1G7B8), Z. tritici strain 3D1

```
Zt_KP6_1     MAPIFTYAVAALAFAQSAYAVVYAARCKFGNPLVQNNRITRAVCDLTNEHTTKDGSWHYV
Zt_SMY22152 MAPIFTYAVAALAFAQSAYAVVYAARCKFGNPLVQNNRITRAVCDLTNEHTTKDGSWHYV
Zt_SMR48303 MAPIFTYAVAALAFAQSAYAVVYAARCKFGNPLVQNNRITRAVCDLTNEHTTKDGSWHYV
Zt_SMR49455 MAPIFTYAVAALAFAQSAYAVVYAARCKFGNPLVQNNRITRAVCDLTNEHTTKDGSWHYV
Zt_SMQ48456 MAPIFTYAVAALAFAQSAYAVVYAARCKFGNPLVQDNRITRAVCNLTNEHTTKDGSWHYV

Zt_KP6_1     EVDNECKYLAGDNPRDQPGWAVFVKYCTYYKGVPDA
Zt_SMY22152 EVDNECKYLAGDNPRDQPGWAVFVKYCTYYKGVPDA
Zt_SMR48303 EVDNECKYLAGDNPRDQPGWAVFVKYCTYYKGVPDA
Zt_SMR49455 EVDNECKYLAGDNPRDQPGWAVFVKYCTYYKGVPDA
Zt_SMQ48456 EVDNECKYLAGDNPRDQPGWAVFVKYCTYYKGYPDA
```

Clustal omega aligment, the amino acids highlighted in red differs from those of SMR48303

Sup. Fig.1-A4 Protein sequence used for heterologous production of Zt-KP6-1

The mature protein without signal peptide was used for protein production in *E. coli.* The N-terminal amino acids (AHM) corresponds to the remains of the protease cleavage site.

```
>mature ZtKP6-1, PDB_6QPK (SMR48303.1, A0A2H1G421) (AHM vector)
(AHM)AVVYAARCKFGNPLVQNNRITRAVCDLTNEHTTKDGSWHYVEVDNECKYLAGDNPRDQPGWAVFVKYCTYYKGVPDA
```

**Sup. Fig.1-B Annotation of IPO323 *Z. tritici* CDS Zt-Mycgr3-106176 (Zt-KP4-1)**

Sup. Fig.1-B1 Annotation of Zt-Mycgr3-106176 (Zt-KP4-1, chr_11:1117942-1119250) using transcript evidence (Lapalu et al., 2023)

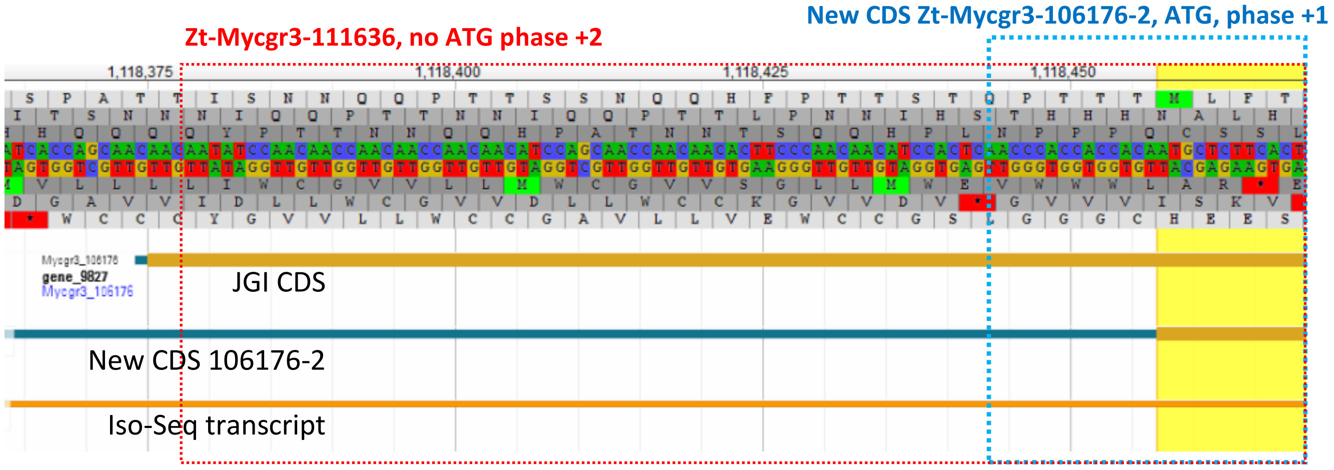


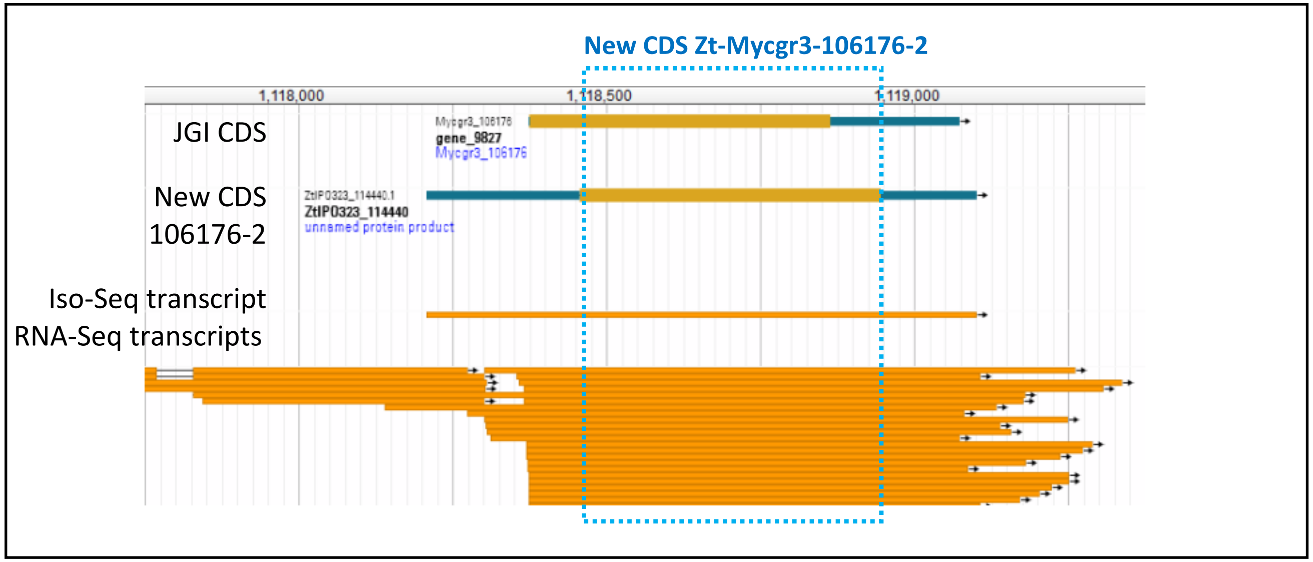

Sup. Fig.1-A2 Protein sequences of Zt-Mycgr3-106176 and Zt-Mycgr3-106176-2

Zt-Mycgr3-106176 and Zt-Mycgr3-106176-2 protein sequences did not share significant homology due to differences in CDS prediction.

```
>Zt-Mycgr3-106176
NNIQQPTTNNIQQPTTLPNNIHSTHHHNALHSKPHPRGPPHHLRPRLSPCPERRRYGRHHRTPQQLRRLHIRPRDRQRRQRPI
QVGLPAAPRRLHRQTEQVLAHLRPSHHWYRSHLPILRDGGCLRYVRLDRTGRHYGSNEGLVELVEGWRNDAGSGRDAGG

>Zt-Mycgr3-106176-2, chr_11:1118208-1119101
MLFTQSLTLVALLTTSALASPLAQNGGGTAGTTGLRNNCDGSTFVPVTGSAGNAPSKWDCQLLRDGYIAKQNKSWLISGPRII
GTVRTCQFSATVDVSGTSGWIGRDDIMDLMKDSLNLWKDGETTQVQGAMQVGESGDVNCVAGKNGEGQKVRIAWTLGHS
```

Sup. Fig.1-B3 Proteins similar to Zt-KP4-1 in annotated *Z. tritici* genomes

SMQ55345.1 (Uniprot A0A1X7S6U3) *Z. tritici* strain 3D7
SMR63670.1 (Uniprot A0A2H1HCW5), *Z. tritici* strain 3D1
SMR60557.1 (Uniprot A0A2H1H404) *Z. tritici* strain 1E4, 8ACX_A
SMY29030.1 (Uniprot A0A1Y6M012) *Z. tritici* strain 1A5
106176-2, (re-annotation, see before), *Z. tritici* strain IPO323
Zt-NIP1, *Z. tritici* strain IPO323 (see below).

```
>Zt_NIP1 IPO323 (Ben M'Barek et al., 2015)
MLFTQSLTLVALLTTSALASPLAQNGGGTAGTTGLRNNCDGSTFVPVTGSAGNAPSKWDCQLLRDGYIAKQNKSWLISGPRII
GTVRTCQFSATVDVSGTSGWIGRDDIMDLMKDSLNLWKDGETTQVQGQMQVGESGDVNCVAGKNGEGQKVRIAWTLGHS
```

Sup. Fig.1-B3 Proteins similar to Zt-KP4-1 in annotated *Z. tritici* genomes

```
SMY29030.1 MLFTQSLTLVALLTTSALASPLAQNGGGTAGTTGLRNNCDGSTFVPVTGSAGNAPSKYDC 60
SMQ55345.1 MLFTQSLTLVALLTTSALASPLAQNGGGTAGTTGLRNNCDGSTFVPVTGSAGNAPSKWDC 60
SMR63670.1 MLFTQSLTLVALLTTSALASPLAQNGGGTAGTTGLRNNCDGSTFVPVTGSAGNAPSKWDC 60
SMR60557.1 MLFTQSLTLVALLTTSALASPLAQNGGGTAGTTGLRNNCDGSTFVPVTGSAGNAPSKWDC 60
Zt_NIP1    MLFTQSLTLVALLTTSALASPLAQNGGGTAGTTGLRNNCDGSTFVPVTGSAGNAPSKWDC 60
106176-2   MLFTQSLTLVALLTTSALASPLAQNGGGTAGTTGLRNNCDGSTFVPVTGSAGNAPSKWDC 60
           *********************************************************:**

SMY29030.1 QLLRDGYIAKQNKSWLISGPRIIGTVRTCQFSATVDVSGTSGWIGRDDIMDLMKDSLNLW 120
SMQ55345.1 QLLRDGYIAKQNKSWLISGPRIIGTVRTCQFSATVDVSGASGWIGRDDIMDLMRDSLNLW 120
SMR63670.1 QLLRDGYIAKQNKSWLISGPRIIGTVRTCQFSATVDVSGTAGWIGRDDIMDLMRDSLNLW 120
SMR60557.1 QLLRDGYIAKQNKSWLISGPRIIGTVRTCQFSATVDVSGTAGWIGRDDIMDLMKDSLNLW 120
Zt_NIP1    QLLRDGYIAKQNKSWLISGPRIIGTVRTCQFSATVDVSGTSGWIGRDDIMDLMKDSLNLW 120
1O6176-2   QLLRDGYIAKQNKSWLISGPRIIGTVRTCQFSATVDVSGTSGWIGRDDIMDLMKDSLNLW 120
           ***************************************::************:******

SMY29030.1 KDGETTQVQGAMQVGESGDVNCVAGKKGEGQKIK--------CSICSFQFNL 164
SMQ55345.1 KDGETTQVQGAMQVGESGDVNCVAGKNGEGKKVPIAWTLGHS---------- 162
SMR63670.1 KDGETTQVQGAMQVGESGDVNCVAGKNGEGKKVPIAWTLGHS---------- 162
SMR60557.1 KDGETTQVQGAMQVGESGDVNCVAGKKGEGQKVRIAWTLGHS---------- 162
Zt_NIP1    KDGETTQVQGQMQVGESGDVNCVAGKNGEGQKVRIAWTLGHS---------- 162
106176-2   KDGETTQVQGAMQVGESGDVNCVAGKNGEGQKVRIAWTLGHS---------- 162
             ********** ***************:***:*:   :     :
```

Clustal omega aligment, the amino acids highlighted in red differs from those of SMR60557.

Sup. Fig.1-B4 Protein sequence used for heterologous production of Zt-KP4-1 in *E. coli*

The mature protein without signal peptide was used for protein production in *E. coli*. The N-terminal amino acid (M) corresponds to the remains of the protease cleavage site.

```
>mature Zt-KP4-1, PDB_8ACX_A (SMR60557.1, A0A2H1H404)(M vector)
(M)SPLAQNGGGTAGTTGLRNNCDGSTFVPVTGSAGNAPSKWDCQLLRDGYIAKQNKSWLISGPRIIGTVRTCQFSATVDVSG
TAGWIGRDDIMDLMKDSLNLWKDGETTQVQGAMQVGESGDVNCVAGKKGEGQKVRIAWTLGHS
```

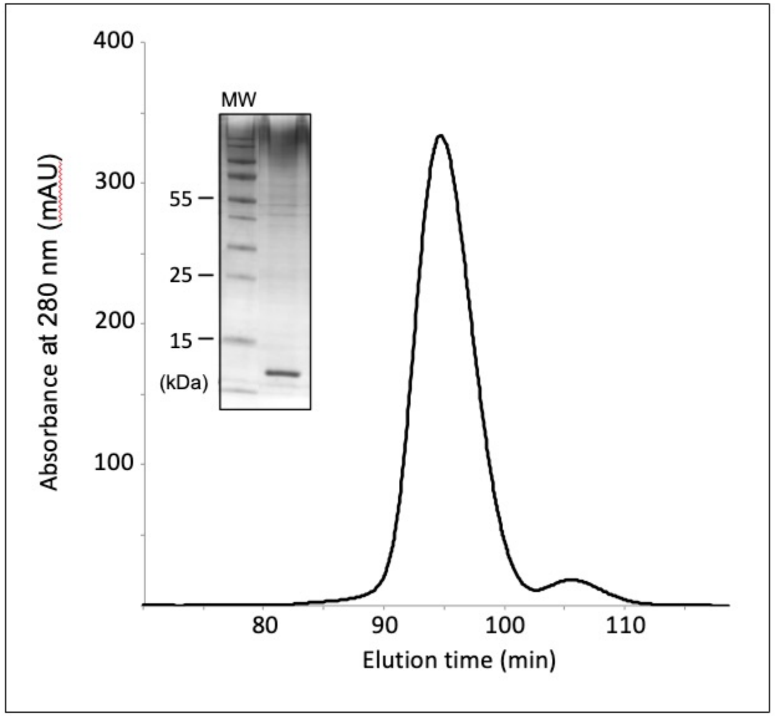


**Supplementary Figure 2. Purification of Zt-KP6-1 by gel filtration**
Gel filtration profile with a Superdex S75 26/60 (GE Healthcare) column at 2.5 ml/min in buffer (20 mM Tris-HCl, pH 8.0, 150 mM NaCl, 1 mM DTT) and SDS-PAGE analysis of purified Zt-KP6-1.
Sizes of the molecular mass markers are indicated on the left of the gel.

**Supplementary Figure 3. NMR analysis of Zt-KP6-1**

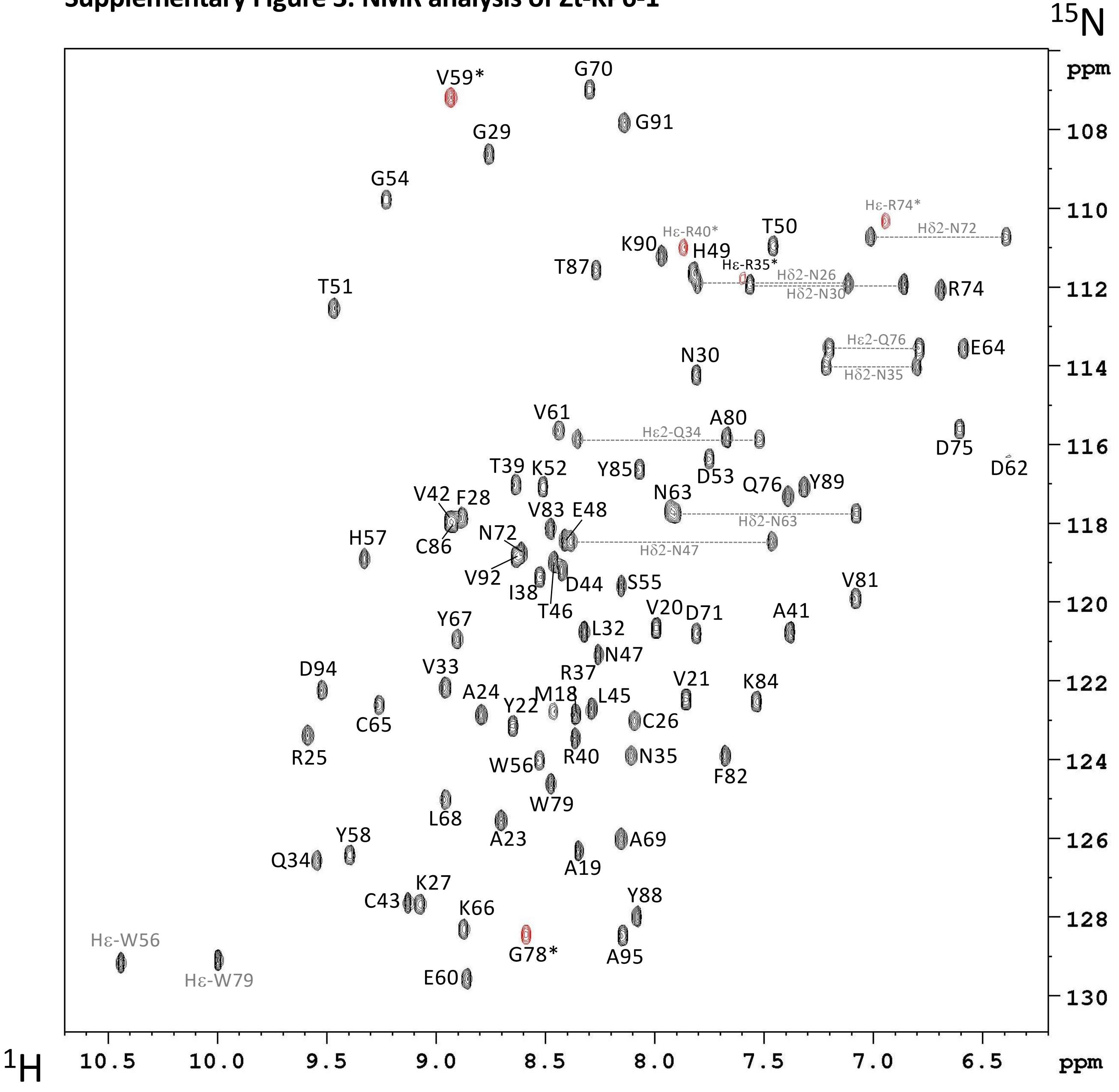


HSQC spectra of Zt-KP6-1 at, 0.7 mM in 20 mM NaCitrate pH 5.4, 150 mM NaCl, 1 mM DTT, 303 K, 800 MHz.
Cross-peak assignments are indicated using one-letter amino acid and number (the asterisk shoiws a folded red peak). Missing residue N36

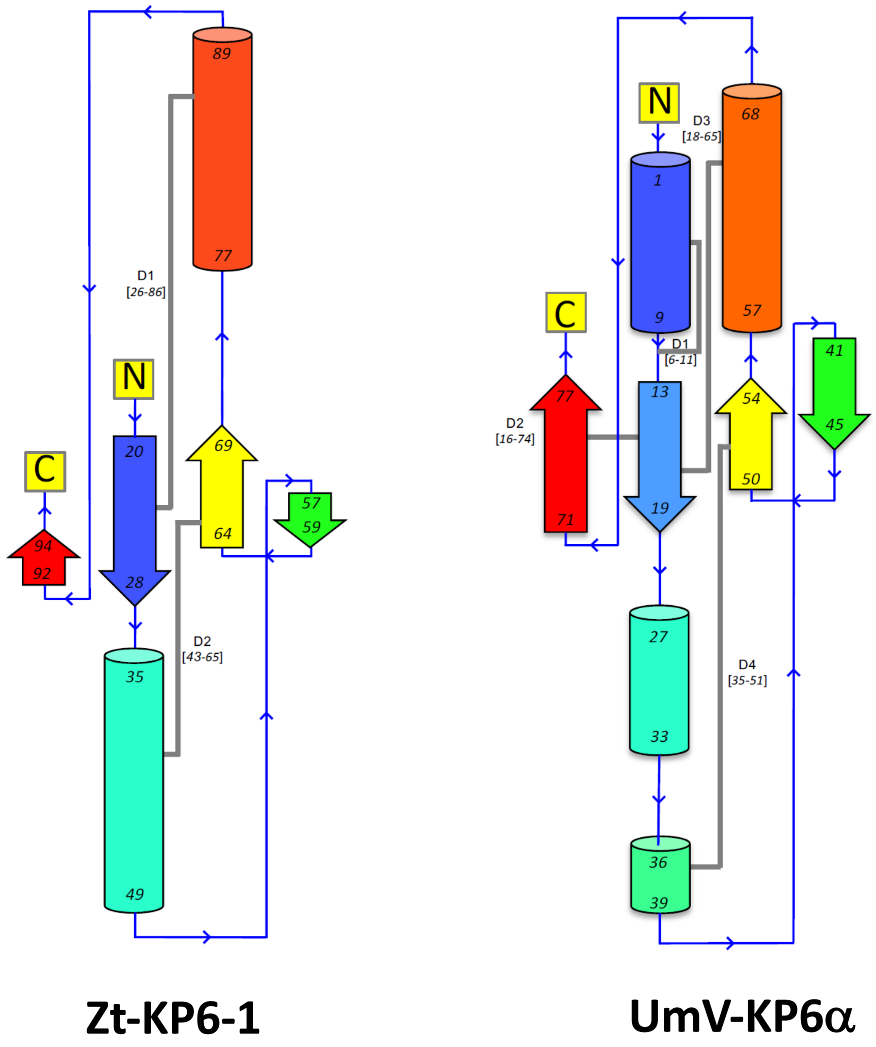


**Supplementary Figure 4. Topology diagrams of Zt-KP6-1 and UmV-KP6α**

The structures were processed by PDBsum and the secondary structure were colored from N-terminus (blue) to C-terminus (red) according to Figure 1. Disulphide Bonds (D) are represented as grey lines.

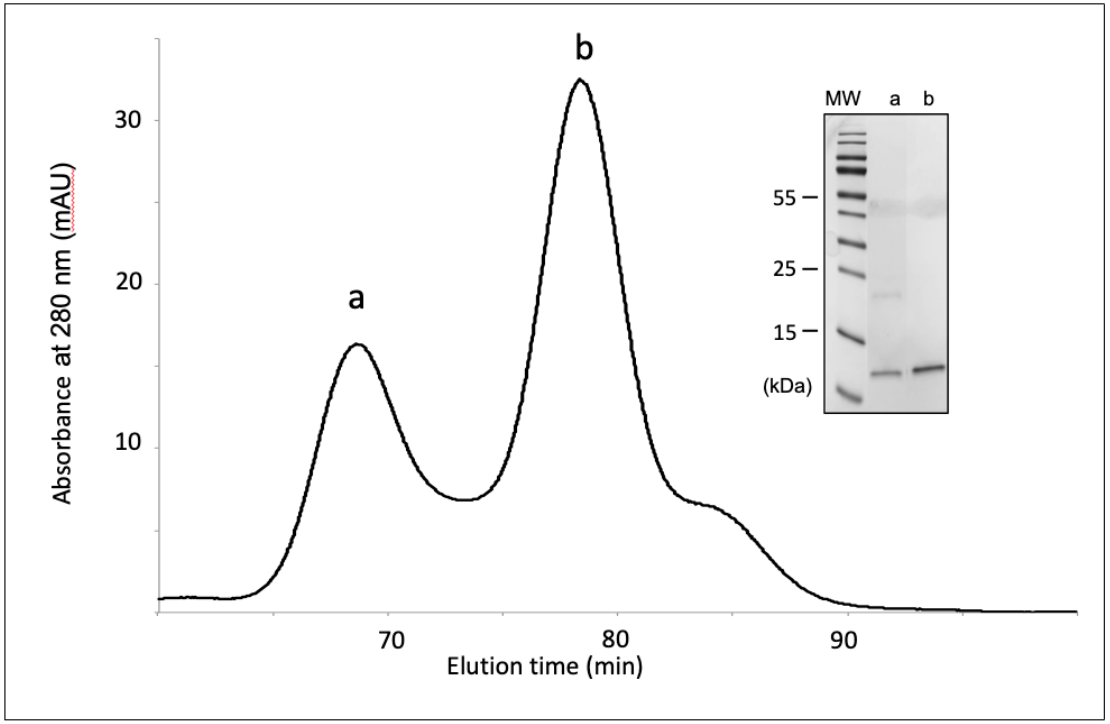


**Supplementary Figure 5. Purification of Zt-KP4-1 by gel filtration**
Gel filtration profile with a Superdex S75 16/60 (GE Healthcare) column at 1 ml/min in buffer (20 mM NaCi, pH 5.6, 150 mM NaCl, 1 mM DTT) and SDS-PAGE analysis of purified Zt-KP4-1. (a) dimer, (b) monomer . Sizes of the molecular mass markers are indicated on the left of the gel.

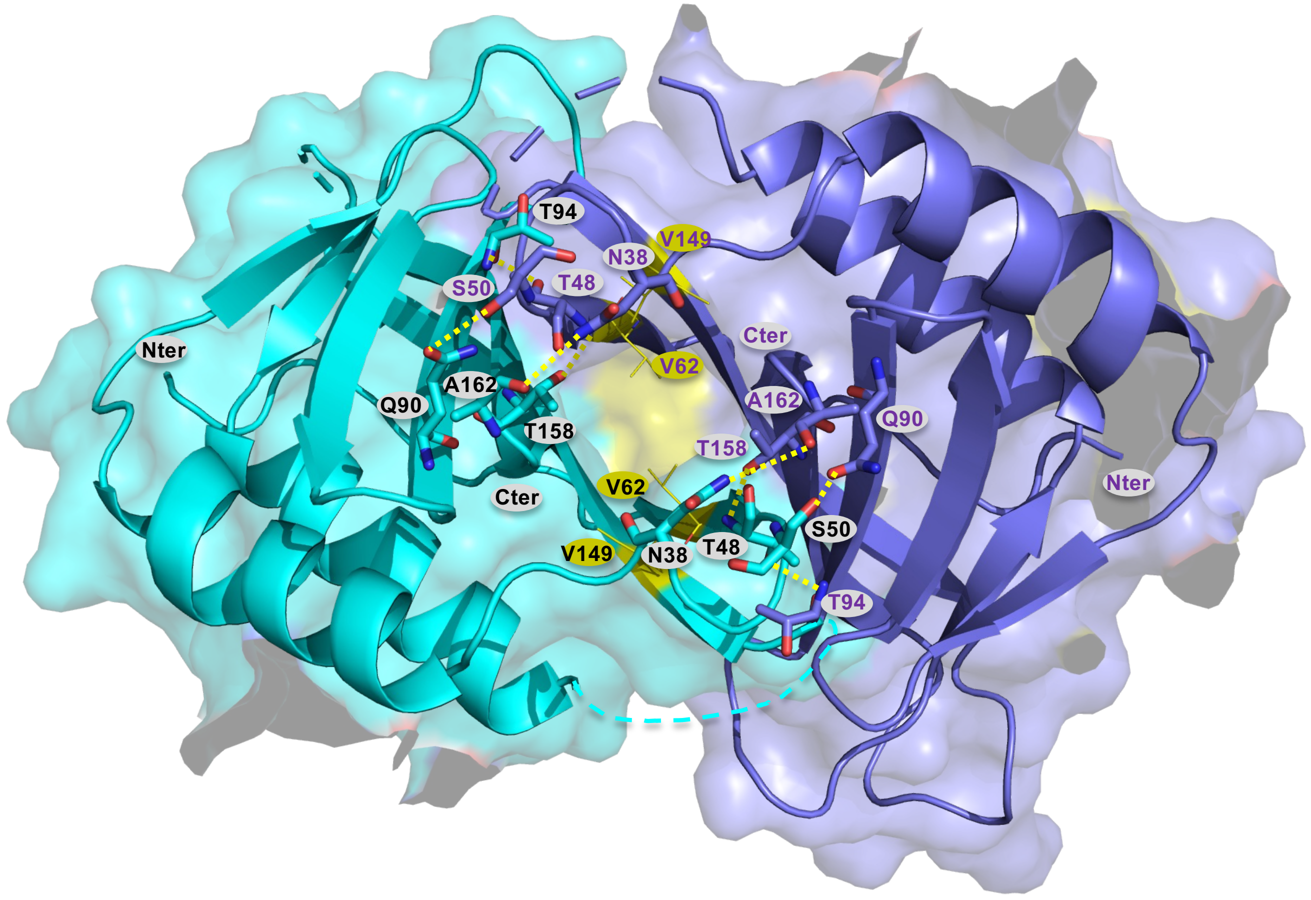


**Supplementary Figure 6. Dimer of Zt-KP4-1 between chains B (cyan) and G (blue).**

Residues involved in hydrogen bonds at the dimer interface are labelled and represented by sticks. Hydrophobics residues involved in the pocket are highlighted in lines and coloured in yellow. According to the GenBank sequence SMR60557.1, the residue numbering must start after the signal peptide at residue S20. T33 in the structure corresponds to T48 in the GenBank sequence.

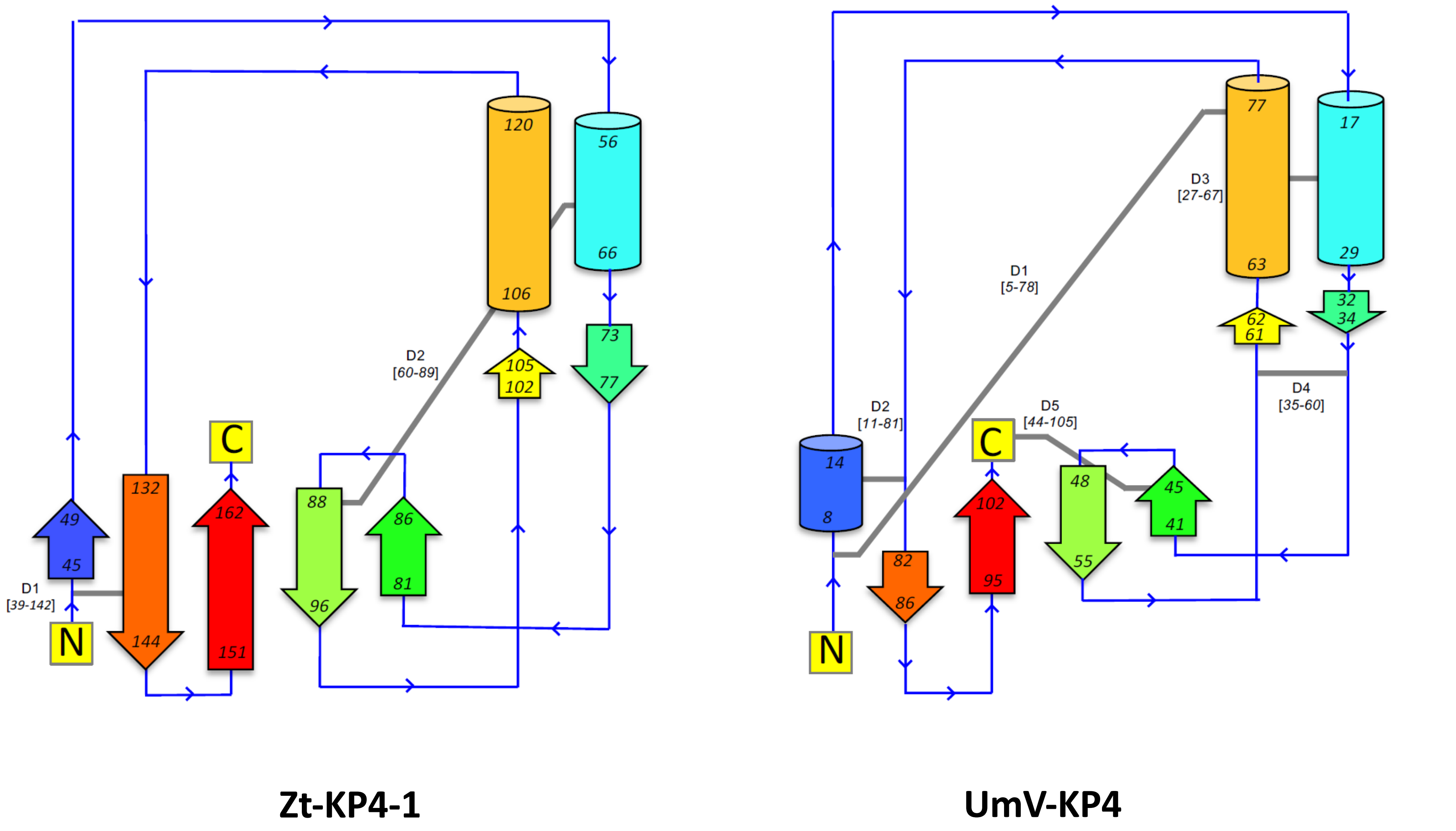


**Supplementary Figure 7.Topology diagrams of Zt-KP4-1 and UmV-KP4**
The structures were processed by PDBsum and the secondary structure were coloured from N-terminus (blue) to C-terminus (red) according to the Figure 2. Disulphide Bonds (D) are represented as grey lines.

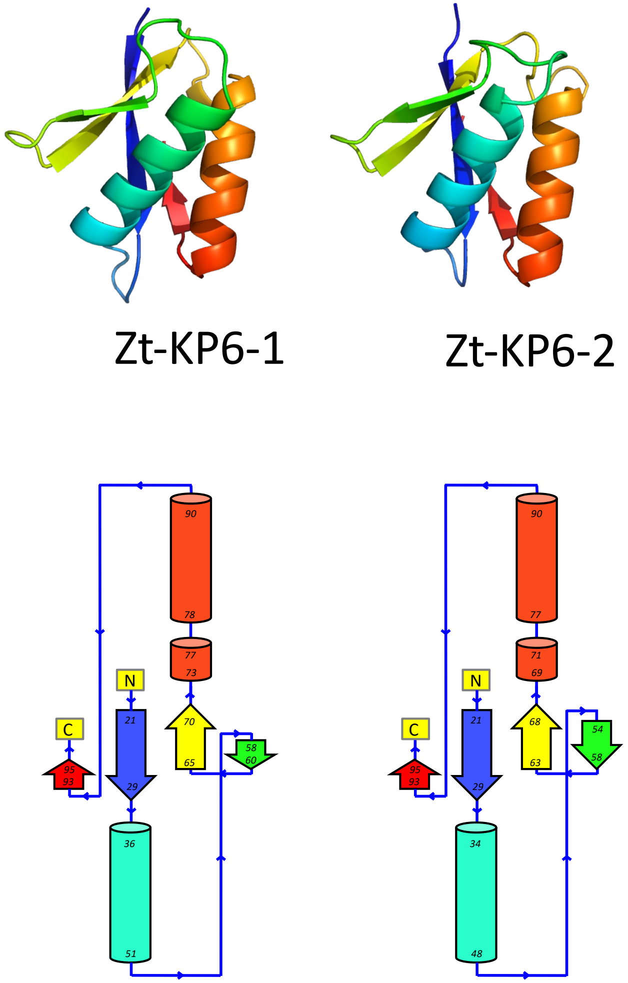


**Supplementary Figure 8: Structure and topology of Zt-KP6-1 and Zt-KP6-2**
Zt-KP6-2 structure was predicted by AlphaFold2. The structures were processed by PDBsum and the secondary structure were coloured from N-terminus (blue) to C-terminus (red) according to Figure 1.

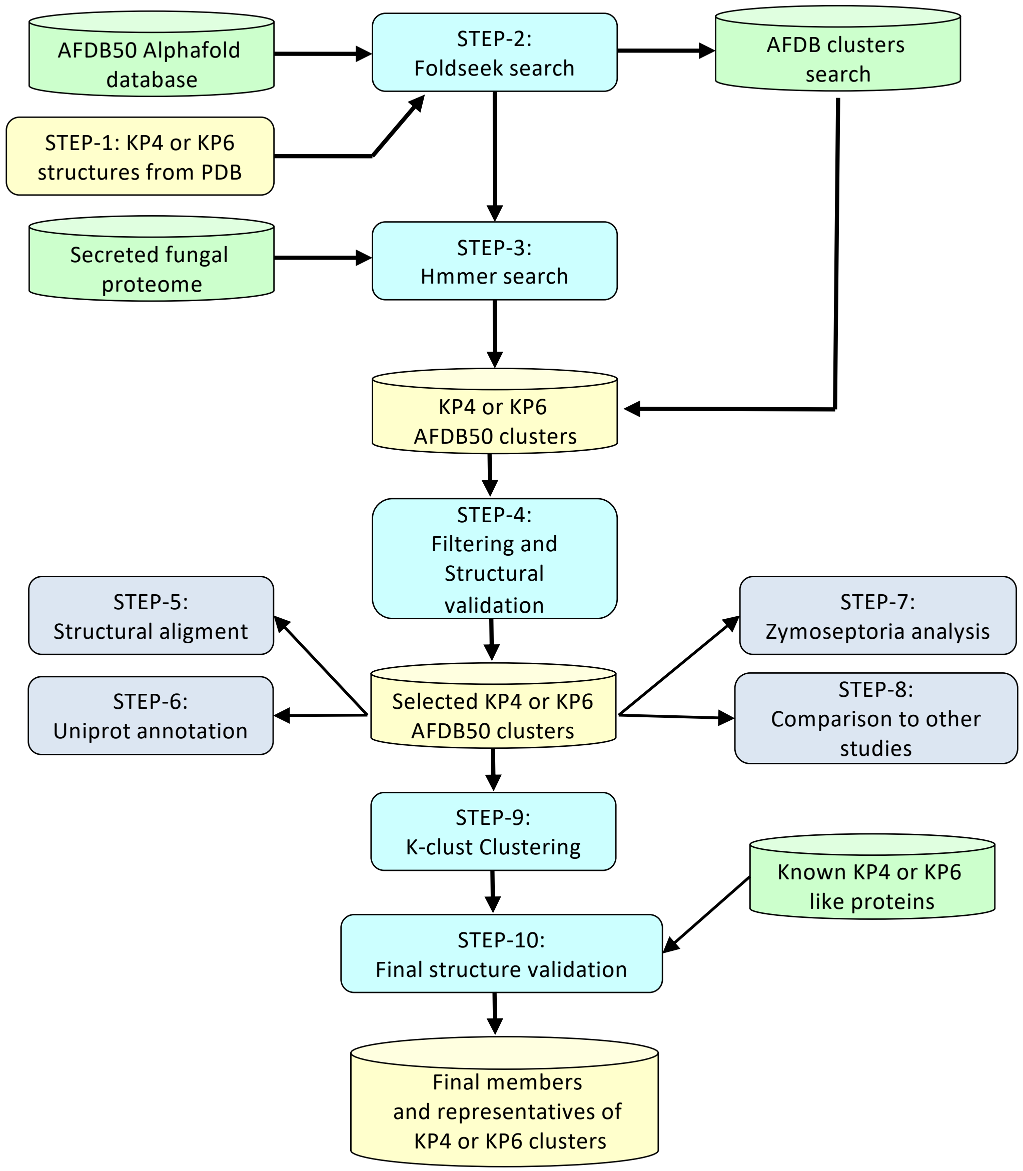


**Supplementary Figure 9: Flow chart of the pipeline used to identify fungal proteins structurally related to Zt-KP6-1 and Zt-KP4-1**

**Supplementary Figure 9:**

- Step 1: Selection of the reference templates (result file1): First, PDB structures were selected as reference templates for the validation of candidate 3D folds (1KPT-A and 8ACX-A in the case of KP4, 4GVB-A and 6QPK-A in the case of KP6).
- Step 2-1: Foldseek search (result files 2 and 4): AlphaFold2 models similar to the reference PDB templates were detected in the AFDB50 database using the Foldseek server (van Kempen et al. 2024).
- Step 2-2: The fungal representatives of the AFDBcluster database (Barrio-Hernandez et al. 2023) were collected (result file 6) if they shared a Foldseek E-value below 1e-3 with any hit from the previous Foldseek search.

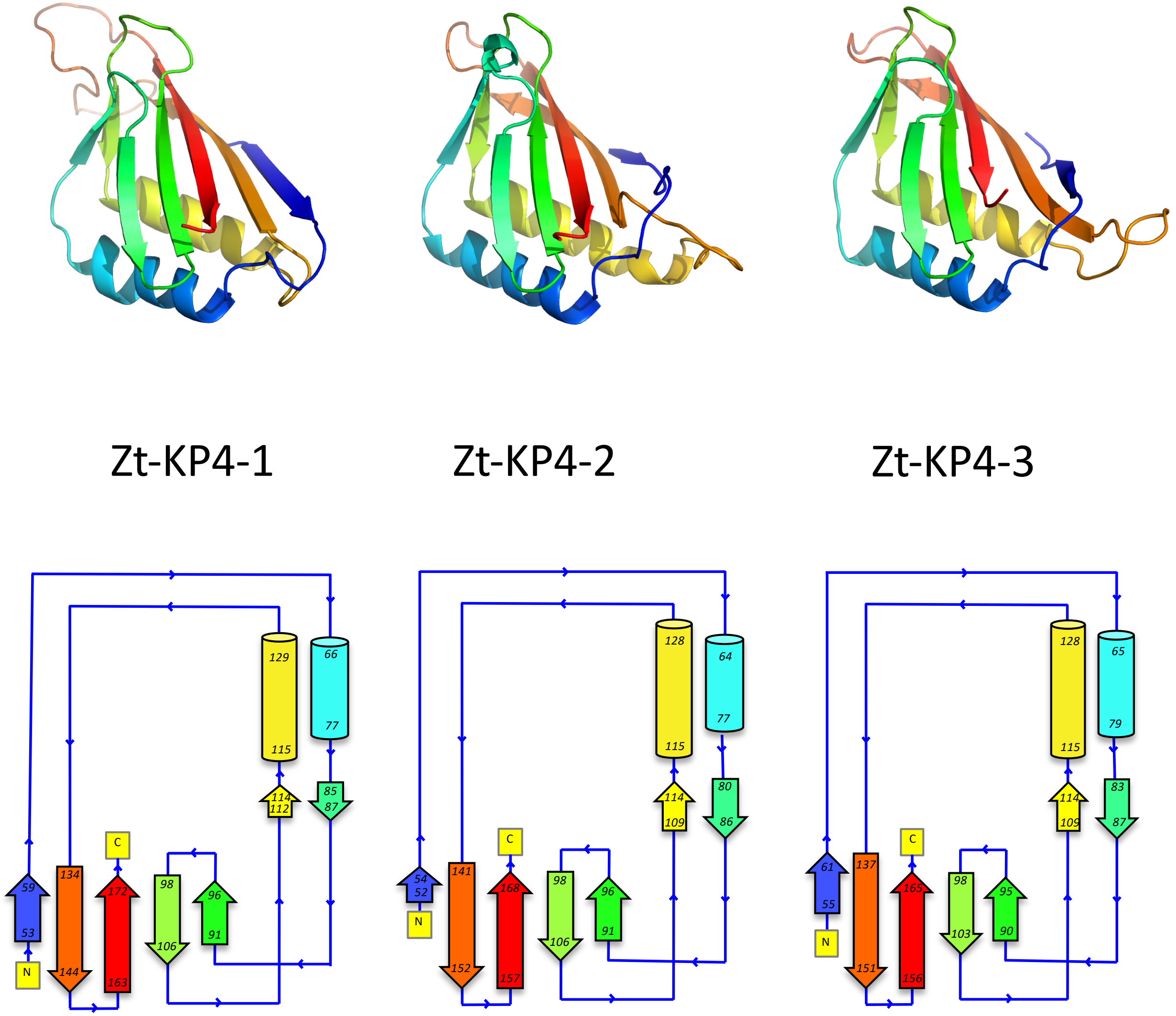


**Supplementary Figure 10: Structure and topology of Zt-KP4-1, Zt-KP4-2 and Zt-KP4-3**
Zt-KP4-2 and Zt-KP4-3 structures were predicted by AlphaFold2. The structures were processed by PDBsum and the secondary structures were coloured from N-terminus (blue) to C-terminus (red) according to Figure 2.

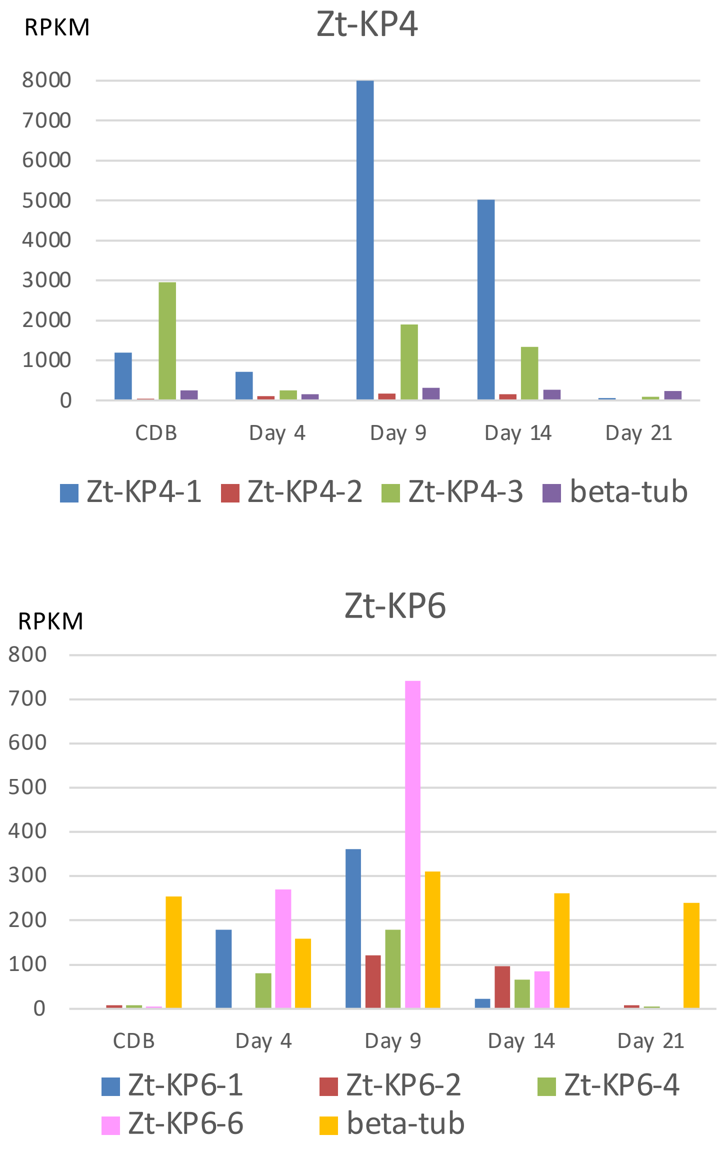


**Supplementary Figure 11: Expression of Zt-KP4-1, Zt-KP4-2, Zt-KP4-3, Zt-KP6-1, Zt-KP6-2, Zt-KP6-4 and Zt-KP6-6 during wheat leaves infection (days after inoculation).**
RNAseq data from Rudd et al. (2015). RPKM (fungal RNAseq **R**eads **P**er **K**ilobase **M**illion)

X-Ray Data collection and refinement statistics

| ***Data collection statistics*** | **Zt-KP6-1** | **Zt-KP4** |
|---|---|---|
| Wavelength (Å) | 0.996 | 0.996 |
| Space group | $P2_12_12_1$ | H3 |
| Cell dimensions | | |
| a, b, c (Å) | 39.39, 57.29, 79.35 | 109.106, 109.106, 252.519 |
| α β γ (°) | 90, 90, 90 | 90, 90, 120 |
| Resolution (Å) | 46.45-1.36 (1.4-1.36) | 75.63 (2.13-1.9) |
| Rmerge | 0.052 (0.9) | 0.088 (1.01) |
| No of reflections | 448377 (15498) | 674238 (43622) |
| No of unique reflections | 33668 | 67360 (5029) |
| I/sI | 24.6 (2.5) | 13 (1.9) |
| Completeness (%) | 91.3 (56.7) | 100 (100) |
| Multiplicity | 12.5 (10.4) | 10 (8.7) |
| ***Refinement*** | | |
| Rwork / Rfree (%) | 0.14/0.183 | 0.175/0.219 |
| RMS (bonds) | 0.017 | 0.015 |
| RMS (angles) | 1.8 | 1.67 |
| Ramachandran favored (%) | 97.95 | 93.59 |
| Ramachandran outliers (%) | 0 | 2.02 |

#Rmerge = Shkl Si |Ii(hkl) - (I(hkl))| / Shkl Si Ii(hkl), where *Ii(hkl)* is the *i*th observation of reflection *hkl* and *<I(hkl)>* is the weighted average intensity for all observations of reflection *hkl*.

**Supplementary Table 1. X-ray Data collection and refinement statistics for Zt-KP6-1 and Zt-KP4-1**

**Supplementary Table 2: NMR and refinement statistics for Zt-KP6-1**

(Zt-KP6-1, 0.7 mM, 20 mM NaCitrate pH 5.4, 150 mM NaCl, 1 mM DTT, 303 K, 800 MHz) PDB:9GWD BMRB ID 34963

| **NMR distance and dihedral constraints** | **Zt-KP6-1** |
|---|---|
| Distance constraints | |
| Total NOE | 1344 |
| Intra-residue | 310 |
| Inter-residue | |
| Sequential ($\|i - j\| = 1$) | 390 |
| Medium-range ($\|i - j\| < 4$) | 236 |
| Long-range ($\|i - j\| > 5$) | 408 |
| Hydrogen bonds | 66 |
| Disulfide bonds | 6 |
| **Structure statistics** | |
| Violations (mean and s.d.) | |
| Max. distance constraint violation (Å) | 0.18 ± 0.03 |
| Deviations from idealized geometry | |
| Bond lengths (Å) | 0.0106 ± 0.0003 |
| Bond angles (º) | 1.1698 ± 0.0431 |
| Impropers (º) | 1.4528 ± 0.0796 |
| **Ramachandran plot (%)** | |
| Most favoured region | 83.2 |
| Additionally allowed region | 13.4 |
| Generously allowed region | 1.9 |
| Disallowed region | 1.5 |
| **Average pairwise *r.m.s.* deviation** (Å)** | |
| Backbone | 0.56 ± 0.12 |
| Heavy | 1.16 ± 0.14 |

** "Pairwise r.m.s. deviation calculated among 20 refined structures for residues 21-95."

**Supplementary Table 3**. **List of the proteins structurally related to Zt-KP6-1 detected using FHAT pipeline.**
https://pat.cbs.cnrs.fr/kp6/fhat_kp6/

**Supplementary Table 4 . List of the proteins structurally related to Zt-KP4-1 detected using FHAT pipeline.**
https://pat.cbs.cnrs.fr/kp4/fhat_kp4/

| KP6 Protein ID | pLDDT | Probability of antimicrobial activity | Prediction | cluster AFDB50 (KclXX-Y) | Pfam domain | Pfam domain name | Group-PCA | Other Name | experimental biological activity |
|---|---|---|---|---|---|---|---|---|---|
| A0A010QJV2_9PEZI | 94,64 | 0,89 | Antimicrobial | Kcl04-C | no | | G1 | Avr-Lm6 like | |
| A0A010RIE2_9PEZI | 92,44 | 0,89 | Antimicrobial | Kcl01-L | no | | G1 | SIX5 like | |
| E5A720_LEPMJ | 95,89 | 0,85 | Antimicrobial | Kcl01-L | no | | G1 | Avr-Lm10A, Lmb_jn3_07875 | |
| E5R515_LEPMJ | 82,99 | 0,93 | Antimicrobial | Kcl04-C | no | | G1 | Avr-Lm6, Lmb_jn3_07862_ | |
| Lmb_jn3_02612 | 76,39 | 0,01 | **Non-antimicrobial** | ND | | | G1 | | |
| Lmb_jn3_08094 | 94,95 | 0,86 | Antimicrobial | ND | | | G1 | | |
| A0A135SXI2_9PEZI | 86,51 | 1,00 | Antimicrobial | Kcl02-G | no | | G2 | | |
| A0A1P8YXM1_PASFU | 90,80 | 0,94 | Antimicrobial | Kcl14-A | no | | G2 | ECP28-2 | |
| A0A1P8YXM5_PASFU | 91,35 | 0,71 | Antimicrobial | Kcl14-A | PF19373 | DUF5948 | G2 | ECP28-3 | |
| A0A1P8YXQ5_PASFU | 86,43 | 0,93 | Antimicrobial | Kcl14-B | PF19373 | DUF5948 | G2 | ECP28-1 | |
| A0A2H1GZA7_ZYMTR | 82,07 | 0,85 | Antimicrobial | Kcl02-D | no | | G2 | Zt-KP6-4 | |
| A0A317AKI4_9PLEO | 83,85 | 0,75 | Antimicrobial | Kcl11-A | no | | G2 | | |
| A0A4V3HRP1_9PEZI | 93,56 | 0,96 | Antimicrobial | Kcl01-C | no | | G2 | | |
| BAS4_MAGO7 | 93,51 | 0,92 | Antimicrobial | Kcl02-G | no | | G2 | BAS4 | |
| F9FAC2_FUSOF | 71,88 | 0,22 | **Non-antimicrobial** | Kcl04-A | no | | G2 | | |
| E5A0W1_LEPMJ | 97,44 | 0,95 | Antimicrobial | Kcl02-G | no | | G2 | Lm-KP6 like, Lmb_jn3_11916 | |
| Lmb_jn3_02094 | 68,18 | 0,24 | **Non-antimicrobial** | ND | | | G2 | | |
| Lmb_jn3_07074 | 90,48 | 0,51 | Antimicrobial | ND | | | G2 | | |
| M1W1H7_CLAP2 | 89,18 | 0,97 | Antimicrobial | Kcl01-C | no | | G2 | | |
| ViKP6fam26 | 88,49 | 0,77 | Antimicrobial | ND | | | G2 | | |
| 4gvbB | na | 0,73 | Antimicrobial | ND | | | G2 | **UmV-KP6-beta** | antifungal |
| 4gvbA | na | 0,90 | Antimicrobial | ND | | | G3 | **UmV-KP6-alpha** | antifungal |
| A0A0F4GGQ1_9PEZI | 73,88 | 0,97 | Antimicrobial | Kcl07-A | no | | G3 | Zb-KP6-6 | |
| A0A0F4GMG5_9PEZI | 78,41 | 0,81 | Antimicrobial | Kcl06-A | no | | G3 | Zb-KP6-2 | |
| A0A2H1FMK1_ZYMTR | 54,75 | 0,82 | Antimicrobial | Kcl07-A | no | | G3 | Zt-KP6-5 | |

| KP6 Protein ID | pLDDT | Probability of antimicrobial activity | Prediction | Pfam domain | Pfam domain name | cluster AFDB50 (KclXX-Y) | PCA | Other Name | Experimental biological activity |
|---|---|---|---|---|---|---|---|---|---|
| A0A010QJV2_9PEZI | 94,64 | 0,89 | Antimicrobial | no | | kcl04-C | G1 | Avr-Lm6 like | |
| A0A010RIE2_9PEZI | 92,44 | 0,89 | Antimicrobial | no | | kcl01-L | G1 | SIX5 like | |
| E5A720_LEPMJ | 95,89 | 0,85 | Antimicrobial | no | | kcl01-L | G1 | Avr-Lm10A, Lmb_jn3_07875 | |
| E5R515_LEPMJ | 82,99 | 0,93 | Antimicrobial | no | | kcl04-C | G1 | Avr-Lm6, Lmb_jn3_07862_ | |
| Lmb_jn3_02612 | 76,39 | 0,01 | **Non-antimicrobial** | | | ND | G1 | | |
| Lmb_jn3_08094 | 94,95 | 0,86 | Antimicrobial | | | ND | G1 | | |
| A0A135SXI2_9PEZI | 86,51 | 1,00 | Antimicrobial | no | | kcl02-G | G2 | | |
| A0A1P8YXM1_PASFU | 90,80 | 0,94 | Antimicrobial | no | | kcl14-A | G2 | ECP28-2 | |
| A0A1P8YXM5_PASFU | 91,35 | 0,71 | Antimicrobial | PF19373 | DUF5948 | kcl14-A | G2 | ECP28-3 | |
| A0A1P8YXQ5_PASFU | 86,43 | 0,93 | Antimicrobial | PF19373 | DUF5948 | kcl14-B | G2 | ECP28-1 | |
| A0A2H1GZA7_ZYMTR | 82,07 | 0,85 | Antimicrobial | no | | kcl02-D | G2 | Zt-KP6-4 | |
| A0A317AKI4_9PLEO | 83,85 | 0,75 | Antimicrobial | no | | kcl11-A | G2 | | |
| A0A4V3HRP1_9PEZI | 93,56 | 0,96 | Antimicrobial | no | | Kcl01-C | G2 | | |
| BAS4_MAGO7 | 93,51 | 0,92 | Antimicrobial | no | | Kcl02-G | G2 | BAS4 | |
| F9FAC2_FUSOF | 71,88 | 0,22 | **Non-antimicrobial** | no | | kcl04-A | G2 | | |
| E5A0W1_LEPMJ | 97,44 | 0,95 | Antimicrobial | no | | kcl02-G | G2 | Lm-KP6 like, Lmb_jn3_11916 | |
| Lmb_jn3_02094 | 68,18 | 0,24 | **Non-antimicrobial** | | | ND | G2 | | |
| Lmb_jn3_07074 | 90,48 | 0,51 | Antimicrobial | | | ND | G2 | | |
| M1W1H7_CLAP2 | 89,18 | 0,97 | Antimicrobial | no | | kcl01-C | G2 | | |
| ViKP6fam26 | 88,49 | 0,77 | Antimicrobial | | | ND | G2 | | |
| 4gvbB | na | 0,73 | Antimicrobial | | | ND | G2 | **UmV-KP6-beta** | antifungal |
| 4gvbA | na | 0,90 | Antimicrobial | | | ND | G3 | **UmV-KP6-alpha** | antifungal |
| A0A0F4GGQ1_9PEZI | 73,88 | 0,97 | Antimicrobial | no | | kcl07-A | G3 | Zb-KP6-6 | |
| A0A0F4GMG5_9PEZI | 78,41 | 0,81 | Antimicrobial | no | | kcl06-A | G3 | Zb-KP6-2 | |
| A0A2H1FMK1_ZYMTR | 54,75 | 0,82 | Antimicrobial | no | | kcl07-A | G3 | Zt-KP6-5 | |
| A0A2H1H0K4_ZYMTR | 77,37 | 0,91 | Antimicrobial | no | | kcl06-A | G3 | Zt-KP6-2 | |
| A0A4R8TM13_9PEZI | 89,81 | 0,83 | Antimicrobial | no | | kcl02-A | G3 | | |
| Lmb_jn3_00833 | 88,06 | 0,91 | Antimicrobial | | | ND | G3 | | |
| Lmb_jn3_06293 | 69,45 | 0,30 | **Non-antimicrobial** | | | ND | G3 | | |
| Lmb_jn3_11952 | 85,84 | 0,77 | Antimicrobial | | | ND | G3 | | |
| ViKP6fam2 | 97,77 | 0,88 | Antimicrobial | | | ND | G3 | | |
| ViKP6fam23 | 80,18 | 0,94 | Antimicrobial | | | ND | G3 | | |
| ViKP6fam5 | 89,01 | 0,98 | Antimicrobial | | | ND | G3 | | |
| A0A2H1G421 | na | 0,77 | Antimicrobial | no | | kcl06-B | G3 | **Zt-KP6-1, 6qpk** | antifungal |
| Lmb_jn3_01852 | 53,92 | 0,66 | Antimicrobial | | | ND | ND | | |

**Supplementary Table 5. Prediction of the antimicrobial activity of fungal KP4 and KP6 proteins using AMAPEC [88].**

**Supplemental File S1: Nucleotide sequence of synthetic genes** (clonning NdeI, BamHI) **and plamid used in this study**

*>Zt-KP6-1 in pET_SB (clonning NdeI, BamHI)*
TTCTTGAAGACGAAAGGGCCTCGTGATACGCCTATTTTTATAGGTTAATGTCATGATAATAATGGTTT
CTTAGACGTCAGGTGGCACTTTTCGGGGAAATGTGCGCGGAACCCCTATTTGTTTATTTTTCTAAATA
CATTCAAATATGTATCCGCTCATGAGACAATAACCCTGATAAATGCTTCAATAATATTGAAAAAGGAA
GAGTATGAGTATTCAACATTTCCGTGTCGCCCTTATTCCCTTTTTTGCGGCATTTTGCCTTCCTGTTTTT
GCTCACCCAGAAACGCTGGTGAAAGTAAAAGATGCTGAAGATCAGTTGGGTGCACGAGTGGGTTAC
ATCGAACTGGATCTCAACAGCGGTAAGATCCTTGAGAGTTTTCGCCCCGAAGAACGTTTTCCAATGAT
GAGCACTTTTAAAGTTCTGCTATGTGGCGCGGTATTATCCCGTGTTGACGCCGGGCAAGAGCAACTC
GGTCGCCGCATACACTATTCTCAGAATGACTTGGTTGAGTACTCACCAGTCACAGAAAAGCATCTTAC
GGATGGCATGACAGTAAGAGAATTATGCAGTGCTGCCATAACCATGAGTGATAACACTGCGGCCAA
CTTACTTCTGACAACGATCGGAGGACCGAAGGAGCTAACCGCTTTTTTGCACAACATGGGGGATCAT
GTAACTCGCCTTGATCGTTGGGAACCGGAGCTGAATGAAGCCATACCAAACGACGAGCGTGACACC
ACGATGCCTGCAGCAATGGCAACAACGTTGCGCAAACTATTAACTGGCGAACTACTTACTCTAGCTTC
CCGGCAACAATTAATAGACTGGATGGAGGCGGATAAAGTTGCAGGACCACTTCTGCGCTCGGCCCTT
CCGGCTGGCTGGTTTATTGCTGATAAATCTGGAGCCGGTGAGCGTGGGTCTCGCGGTATCATTGCAG
CACTGGGGCCAGATGGTAAGCCCTCCCGTATCGTAGTTATCTACACGACGGGGAGTCAGGCAACTAT
GGATGAACGAAATAGACAGATCGCTGAGATAGGTGCCTCACTGATTAAGCATTGGTAACTGTCAGA
CCAAGTTTACTCATATATACTTTAGATTGATTTAAAACTTCATTTTTAATTTAAAAGGATCTAGGTGAA
GATCCTTTTTGATAATCTCATGACCAAAATCCCTTAACGTGAGTTTTCGTTCCACTGAGCGTCAGACCC
CGTAGAAAAGATCAAAGGATCTTCTTGAGATCCTTTTTTTCTGCGCGTAATCTGCTGCTTGCAAACAA
AAAAACCACCGCTACCAGCGGTGGTTTGTTTGCCGGATCAAGAGCTACCAACTCTTTTTCCGAAGGTA
ACTGGCTTCAGCAGAGCGCAGATACCAAATACTGTCCTTCTAGTGTAGCCGTAGTTAGGCCACCACTT
CAAGAACTCTGTAGCACCGCCTACATACCTCGCTCTGCTAATCCTGTTACCAGTGGCTGCTGCCAGTG
GCGATAAGTCGTGTCTTACCGGGTTGGACTCAAGACGATAGTTACCGGATAAGGCGCAGCGGTCGG
GCTGAACGGGGGGTTCGTGCACACAGCCCAGCTTGGAGCGAACGACCTACACCGAACTGAGATACC
TACAGCGTGAGCTATGAGAAAGCGCCACGCTTCCCGAAGGGAGAAAGGCGGACAGGTATCCGGTA
AGCGGCAGGGTCGGAACAGGAGAGCGCACGAGGGAGCTTCCAGGGGGAAACGCCTGGTATCTTTA
TAGTCCTGTCGGGTTTCGCCACCTCTGACTTGAGCGTCGATTTTTGTGATGCTCGTCAGGGGGGCGG
AGCCTATGGAAAAACGCCAGCAACGCGGCCTTTTTACGGTTCCTGGCCTTTTGCTGGCCTTTTGCTCA
CATGTTCTTTCCTGCGTTATCCCCTGATTCTGTGGATAACCGTATTACCGCCTTTGAGTGAGCTGATAC
CGCTCGCCGCAGCCGAACGACCGAGCGCAGCGAGTCAGTGAGCGAGGAAGCGGAAGAGCGCCTGA
TGCGGTATTTTCTCCTTACGCATCTGTGCGGTATTTCACACCGCATATATGGTGCACTCTCAGTACAAT
CTGCTCTGATGCCGCATAGTTAAGCCAGTATACACTCCGCTATCGCTACGTGACTGGGTCATGGCTGC
GCCCCGACACCCGCCAACACCCGCTGACGCGCCCTGACGGGCTTGTCTGCTCCCGGCATCCGCTTAC
AGACAAGCTGTGACCGTCTCCGGGAGCTGCATGTGTCAGAGGTTTTCACCGTCATCACCGAAACGCG
CGAGGCAGCTGCGGTAAAGCTCATCAGCGTGGTCGTGAAGCGATTCACAGATGTCTGCCTGTTCATC
CGCGTCCAGCTCGTTGAGTTTCTCCAGAAGCGTTAATGTCTGGCTTCTGATAAAGCGGGCCATGTTAA
GGGCGGTTTTTTCCTGTTTGGTCACTGATGCCTCCGTGTAAGGGGGATTTCTGTTCATGGGGGTAAT
GATACCGATGAAACGAGAGAGGATGCTCACGATACGGGTTACTGATGATGAACATGCCCGGTTACT
GGAACGTTGTGAGGGTAAACAACTGGCGGTATGGATGCGGCGGGACCAGAGAAAAATCACTCAGG
GTCAATGCCAGCGCTTCGTTAATACAGATGTAGGTGTTCCACAGGGTAGCCAGCAGCATCCTGCGAT
GCAGATCCGGAACATAATGGTGCAGGGCGCTGACTTCCGCGTTTCCAGACTTTACGAAACACGGAAA
CCGAAGACCATTCATGTTGTTGCTCAGGTCGCAGACGTTTTGCAGCAGCAGTCGCTTCACGTTCGCTC
GCGTATCGGTGATTCATTCTGCTAACCAGTAAGGCAACCCCGCCAGCCTAGCCGGGTCCTCAACGAC

AGGAGCACGATCATGCGCACCCGTGGCCAGGACCCAACGCTGCCCGAGATGCGCCGCGTGCGGCTG
CTGGAGATGGCGGACGCGATGGATATGTTCTGCCAAGGGTTGGTTTGCGCATTCACAGTTCTCCGCA
AGAATTGATTGGCTCCAATTCTTGGAGTGGTGAATCCGTTAGCGAGGTGCCGCCGGCTTCCATTCAG
GTCGAGGTGGCCCGGCTCCATGCACCGCGACGCAACGCGGGGAGGCAGACAAGGTATAGGGCGGC
GCCTACAATCCATGCCAACCCGTTCCATGTGCTCGCCGAGGCGGCATAAATCGCCGTGACGATCAGC
GGTCCAGTGATCGAAGTTAGGCTGGTAAGAGCCGCGAGCGATCCTTGAAGCTGTCCCTGATGGTCG
TCATCTACCTGCCTGGACAGCATGGCCTGCAACGCGGGCATCCCGATGCCGCCGGAAGCGAGAAGA
ATCATAATGGGGAAGGCCATCCAGCCTCGCGTCGCGAACGCCAGCAAGACGTAGCCCAGCGCGTCG
GCCGCCATGCCGGCGATAATGGCCTGCTTCTCGCCGAAACGTTTGGTGGCGGGACCAGTGACGAAG
GCTTGAGCGAGGGCGTGCAAGATTCCGAATACCGCAAGCGACAGGCCGATCATCGTCGCGCTCCAG
CGAAAGCGGTCCTCGCCGAAAATGACCCAGAGCGCTGCCGGCACCTGTCCTACGAGTTGCATGATAA
AGAAGACAGTCATAAGTGCGGCGACGATAGTCATGCCCCGCGCCCACCGGAAGGAGCTGACTGGGT
TGAAGGCTCTCAAGGGCATCGGTCGAGATCCCGGTGCCTAATGAGTGAGCTAACTTACATTAATTGC
GTTGCGCTCACTGCCCGCTTTCCAGTCGGGAAACCTGTCGTGCCAGCTGCATTAATGAATCGGCCAAC
GCGCGGGGAGAGGCGGTTTGCGTATTGGGCGCCAGGGTGGTTTTTCTTTTCACCAGTGAGACGGGC
AACAGCTGATTGCCCTTCACCGCCTGGCCCTGAGAGAGTTGCAGCAAGCGGTCCACGCTGGTTTGCC
CCAGCAGGCGAAAATCCTGTTTGATGGTGGTTAACGGCGGGATATAACATGAGCTGTCTTCGGTATC
GTCGTATCCCACTACCGAGATATCCGCACCAACGCGCAGCCCGGACTCGGTAATGGCGCGCATTGCG
CCCAGCGCCATCTGATCGTTGGCAACCAGCATCGCAGTGGGAACGATGCCCTCATTCAGCATTTGCA
TGGTTTGTTGAAAACCGGACATGGCACTCCAGTCGCCTTCCCGTTCCGCTATCGGCTGAATTTGATTG
CGAGTGAGATATTTATGCCAGCCAGCCAGACGCAGACGCGCCGAGACAGAACTTAATGGGCCCGCT
AACAGCGCGATTTGCTGGTGACCCAATGCGACCAGATGCTCCACGCCCAGTCGCGTACCGTCTTCAT
GGGAGAAAATAATACTGTTGATGGGTGTCTGGTCAGAGACATCAAGAAATAACGCCGGAACATTAG
TGCAGGCAGCTTCCACAGCAATGGCATCCTGGTCATCCAGCGGATAGTTAATGATCAGCCCACTGAC
GCGTTGCGCGAGAAGATTGTGCACCGCCGCTTTACAGGCTTCGACGCCGCTTCGTTCTACCATCGAC
ACCACCACGCTGGCACCCAGTTGATCGGCGCGAGATTTAATCGCCGCGACAATTTGCGACGGCGCGT
GCAGGGCCAGACTGGAGGTGGCAACGCCAATCAGCAACGACTGTTTGCCCGCCAGTTGTTGTGCCA
CGCGGTTGGGAATGTAATTCAGCTCCGCCATCGCCGCTTCCACTTTTTCCCGCGTTTTCGCAGAAACG
TGGCTGGCCTGGTTCACCACGCGGGAAACGGTCTGATAAGAGACACCGGCATACTCTGCGACATCGT
ATAACGTTACTGGTTTCACATTCACCACCCTGAATTGACTCTCTTCCGGGCGCTATCATGCCATACCGC
GAAAGGTTTTGCGCCATTCGATGGTGTCCGGGATCTCGACGCTCTCCCTTATGCGACTCCTGCATTAG
GAAGCAGCCCAGTAGTAGGTTGAGGCCGTTGAGCACCGCCGCCGCAAGGAATGGTGCATGCAAGG
AGATGGCGCCCAACAGTCCCCCGGCCACGGGGCCTGCCACCATACCCACGCCGAAACAAGCGCTCAT
GAGCCCGAAGTGGCGAGCCCGATCTTCCCCATCGGTGATGTCGGCGATATAGGCGCCAGCAACCGC
ACCTGTGGCGCCGGTGATGCCGGCCACGATGCGTCCGGCGTAGAGGATCGAGATCTCGATCCCGCG
AAATTAATACGACTCACTATAGGGGAATTGTGAGCGGATAACAATTCCCCTCTAGAAATAATTTTGTT
TAACTTTAAGAAGGAGATATACCATGAAAAAGACAGCTATCGCGATTGCAGTGGCACTGGCTGGTTT
CGCTACCGTAGCGCAGGCCGCTCCGCAAGATAACACTAGCATGGGCAGCAGCCATCATCATCATCAT
CACAGCAGCGGCAGAGAAAACTTGTATTTCCAGGGCCATATGGCTGTCGTTTATGCCGCACGCTGCA
AATTTGGCAATCCGTTAGTGCAGAACAACCGCATTACTCGTGCCGTATGTGACCTGACAAACGAACA
TACCACCAAAGATGGTAGCTGGCACTATGTGGAAGTCGACAATGAGTGCAAATATCTGGCTGGCGA
TAATCCGCGTGATCAACCTGGTTGGGCGGTATTCGTGAAGTACTGTACGTACTACAAAGGGGTTCCA
GATGCGTAGGGATCCTAATAACTAAGTAAACTAGTGCCTGAGCAATAACTAGCATAACCCCCTTGGG
GCCTCTAAACGGGTCTTGAGGGGTTTTTTGCTGAAAGGAGGAACTATATCCGGATATCCCGCAAGAG
GCCCGGCAGTACCGGCATAACCAAGCCTATGCCTACAGCATCCAGGGTGACGGTGCCGAGGATGAC
GATGAGCGCATTGTTAGATTTCATACACGGTGCCTGACTGCGTTAGCAATTTAACTGTGATAAACTAC
CGCATTAAAGCTTATCGATGATAAGCTGTCAAACATGAGAA

>Zt-KP4 in pDb_ccdb_pepL_his_3C *(clonning Nde1 Xho)*

TCCAGGGGCCCCATATGTCTCCGCTGGCTCAAAACGGTGGCGGAACAGCAGGCACAACAGGCTTAC
GCAATAATTGTGACGGCTCTACATTCGTCCCAGTTACGGGTTCCGCTGGAAACGCGCCTAGCAAATG
GGACTGTCAGTTGTTACGTGATGGCTACATCGCTAAACAAAACAAGAGTTGGTTAATTAGCGGTCCA
CGCATCATTGGGACTGTTCGCACTTGTCAGTTCAGTGCGACAGTCGACGTTTCCGGAACGGCTGGGT
GGATTGGCCGCGACGACATCATGGACTTAATGAAAGACTCGTTAAACTTATGGAAGGACGGAGAGA
CGACACAGGTACAAGGGGCTATGCAGGTAGGAGAAAGTGGCGACGTGAATTGTGTGGCAGGTAAA
AAGGGGGAAGGTCAGAAAGTCCGCATCGCGTGGACATTGGGCCATTCCTAATGACTCGAGCACCAC
CACC

> pDb_ccdb_pepL_his_3C
TGGCGAATGGGACGCGCCCTGTAGCGGCGCATTAAGCGCGGCGGGTGTGGTGGTTACGCGCAGCG
TGACCGCTACACTTGCCAGCGCCCTAGCGCCCGCTCCTTTCGCTTTCTTCCCTTCCTTTCTCGCCACGTT
CGCCGGCTTTCCCCGTCAAGCTCTAAATCGGGGGCTCCCTTTAGGGTTCCGATTTAGTGCTTTACGGC
ACCTCGACCCCAAAAAACTTGATTAGGGTGATGGTTCACGTAGTGGGCCATCGCCCTGATAGACGGT
TTTTCGCCCTTTGACGTTGGAGTCCACGTTCTTTAATAGTGGACTCTTGTTCCAAACTGGAACAACACT
CAACCCTATCTCGGTCTATTCTTTTGATTTATAAGGGATTTTGCCGATTTCGGCCTATTGGTTAAAAAA
TGAGCTGATTTAACAAAAATTTAACGCGAATTTTAACAAAATATTAACGTTTACAATTTCAGGTGGCA
CTTTTCGGGGAAATGTGCGCGGAACCCCTATTTGTTTATTTTTCTAAATACATTCAAATATGTATCCGC
TCATGAATTAATTCTTAGAAAAACTCATCGAGCATCAAATGAAACTGCAATTTATTCATATCAGGATT
ATCAATACCATATTTTTGAAAAAGCCGTTTCTGTAATGAAGGAGAAAACTCACCGAGGCAGTTCCATA
GGATGGCAAGATCCTGGTATCGGTCTGCGATTCCGACTCGTCCAACATCAATACAACCTATTAATTTC
CCCTCGTCAAAAATAAGGTTATCAAGTGAGAAATCACCATGAGTGACGACTGAATCCGGTGAGAATG
GCAAAAGTTTATGCATTTCTTTCCAGACTTGTTCAACAGGCCAGCCATTACGCTCGTCATCAAAATCAC
TCGCATCAACCAAACCGTTATTCATTCGTGATTGCGCCTGAGCGAGACGAAATACGCGATCGCTGTTA
AAAGGACAATTACAAACAGGAATCGAATGCAACCGGCGCAGGAACACTGCCAGCGCATCAACAATA
TTTTCACCTGAATCAGGATATTCTTCTAATACCTGGAATGCTGTTTTCCCGGGGATCGCAGTGGTGAG
TAACCATGCATCATCAGGAGTACGGATAAAATGCTTGATGGTCGGAAGAGGCATAAATTCCGTCAGC
CAGTTTAGTCTGACCATCTCATCTGTAACATCATTGGCAACGCTACCTTTGCCATGTTTCAGAAACAAC
TCTGGCGCATCGGGCTTCCCATACAATCGATAGATTGTCGCACCTGATTGCCCGACATTATCGCGAGC
CCATTTATACCCATATAAATCAGCATCCATGTTGGAATTTAATCGCGGCCTAGAGCAAGACGTTTCCC
GTTGAATATGGCTCATAACACCCCTTGTATTACTGTTTATGTAAGCAGACAGTTTTATTGTTCATGACC
AAAATCCCTTAACGTGAGTTTTCGTTCCACTGAGCGTCAGACCCCGTAGAAAAGATCAAAGGATCTTC
TTGAGATCCTTTTTTTCTGCGCGTAATCTGCTGCTTGCAAACAAAAAAACCACCGCTACCAGCGGTGG
TTTGTTTGCCGGATCAAGAGCTACCAACTCTTTTTCCGAAGGTAACTGGCTTCAGCAGAGCGCAGATA
CCAAATACTGTCCTTCTAGTGTAGCCGTAGTTAGGCCACCACTTCAAGAACTCTGTAGCACCGCCTAC
ATACCTCGCTCTGCTAATCCTGTTACCAGTGGCTGCTGCCAGTGGCGATAAGTCGTGTCTTACCGGGT
TGGACTCAAGACGATAGTTACCGGATAAGGCGCAGCGGTCGGGCTGAACGGGGGGTTCGTGCACA
CAGCCCAGCTTGGAGCGAACGACCTACACCGAACTGAGATACCTACAGCGTGAGCTATGAGAAAGC
GCCACGCTTCCCGAAGGGAGAAAGGCGGACAGGTATCCGGTAAGCGGCAGGGTCGGAACAGGAGA
GCGCACGAGGGAGCTTCCAGGGGGAAACGCCTGGTATCTTTATAGTCCTGTCGGGTTTCGCCACCTC
TGACTTGAGCGTCGATTTTTGTGATGCTCGTCAGGGGGGCGGAGCCTATGGAAAAACGCCAGCAAC
GCGGCCTTTTTACGGTTCCTGGCCTTTTGCTGGCCTTTTGCTCACATGTTCTTTCCTGCGTTATCCCCTG
ATTCTGTGGATAACCGTATTACCGCCTTTGAGTGAGCTGATACCGCTCGCCGCAGCCGAACGACCGA
GCGCAGCGAGTCAGTGAGCGAGGAAGCGGAAGAGCGCCTGATGCGGTATTTTCTCCTTACGCATCT
GTGCGGTATTTCACACCGCATATATGGTGCACTCTCAGTACAATCTGCTCTGATGCCGCATAGTTAAG

CCAGTATACACTCCGCTATCGCTACGTGACTGGGTCATGGCTGCGCCCCGACACCCGCCAACACCCG
CTGACGCGCCCTGACGGGCTTGTCTGCTCCCGGCATCCGCTTACAGACAAGCTGTGACCGTCTCCGG
GAGCTGCATGTGTCAGAGGTTTTCACCGTCATCACCGAAACGCGCGAGGCAGCTGCGGTAAAGCTC
ATCAGCGTGGTCGTGAAGCGATTCACAGATGTCTGCCTGTTCATCCGCGTCCAGCTCGTTGAGTTTCT
CCAGAAGCGTTAATGTCTGGCTTCTGATAAAGCGGGCCATGTTAAGGGCGGTTTTTTCCTGTTTGGTC
ACTGATGCCTCCGTGTAAGGGGGATTTCTGTTCATGGGGGTAATGATACCGATGAAACGAGAGAGG
ATGCTCACGATACGGGTTACTGATGATGAACATGCCCGGTTACTGGAACGTTGTGAGGGTAAACAAC
TGGCGGTATGGATGCGGCGGGACCAGAGAAAAATCACTCAGGGTCAATGCCAGCGCTTCGTTAATA
CAGATGTAGGTGTTCCACAGGGTAGCCAGCAGCATCCTGCGATGCAGATCCGGAACATAATGGTGC
AGGGCGCTGACTTCCGCGTTTCCAGACTTTACGAAACACGGAAACCGAAGACCATTCATGTTGTTGC
TCAGGTCGCAGACGTTTTGCAGCAGCAGTCGCTTCACGTTCGCTCGCGTATCGGTGATTCATTCTGCT
AACCAGTAAGGCAACCCCGCCAGCCTAGCCGGGTCCTCAACGACAGGAGCACGATCATGCGCACCC
GTGGGGCCGCCATGCCGGCGATAATGGCCTGCTTCTCGCCGAAACGTTTGGTGGCGGGACCAGTGA
CGAAGGCTTGAGCGAGGGCGTGCAAGATTCCGAATACCGCAAGCGACAGGCCGATCATCGTCGCGC
TCCAGCGAAAGCGGTCCTCGCCGAAAATGACCCAGAGCGCTGCCGGCACCTGTCCTACGAGTTGCAT
GATAAAGAAGACAGTCATAAGTGCGGCGACGATAGTCATGCCCCGCGCCCACCGGAAGGAGCTGAC
TGGGTTGAAGGCTCTCAAGGGCATCGGTCGAGATCCCGGTGCCTAATGAGTGAGCTAACTTACATTA
ATTGCGTTGCGCTCACTGCCCGCTTTCCAGTCGGGAAACCTGTCGTGCCAGCTGCATTAATGAATCGG
CCAACGCGCGGGGAGAGGCGGTTTGCGTATTGGGCGCCAGGGTGGTTTTTCTTTTCACCAGTGAGA
CGGGCAACAGCTGATTGCCCTTCACCGCCTGGCCCTGAGAGAGTTGCAGCAAGCGGTCCACGCTGGT
TTGCCCCAGCAGGCGAAAATCCTGTTTGATGGTGGTTAACGGCGGGATATAACATGAGCTGTCTTCG
GTATCGTCGTATCCCACTACCGAGATATCCGCACCAACGCGCAGCCCGGACTCGGTAATGGCGCGCA
TTGCGCCCAGCGCCATCTGATCGTTGGCAACCAGCATCGCAGTGGGAACGATGCCCTCATTCAGCAT
TTGCATGGTTTGTTGAAAACCGGACATGGCACTCCAGTCGCCTTCCCGTTCCGCTATCGGCTGAATTT
GATTGCGAGTGAGATATTTATGCCAGCCAGCCAGACGCAGACGCGCCGAGACAGAACTTAATGGGC
CCGCTAACAGCGCGATTTGCTGGTGACCCAATGCGACCAGATGCTCCACGCCCAGTCGCGTACCGTC
TTCATGGGAGAAAATAATACTGTTGATGGGTGTCTGGTCAGAGACATCAAGAAATAACGCCGGAAC
ATTAGTGCAGGCAGCTTCCACAGCAATGGCATCCTGGTCATCCAGCGGATAGTTAATGATCAGCCCA
CTGACGCGTTGCGCGAGAAGATTGTGCACCGCCGCTTTACAGGCTTCGACGCCGCTTCGTTCTACCAT
CGACACCACCACGCTGGCACCCAGTTGATCGGCGCGAGATTTAATCGCCGCGACAATTTGCGACGGC
GCGTGCAGGGCCAGACTGGAGGTGGCAACGCCAATCAGCAACGACTGTTTGCCCGCCAGTTGTTGT
GCCACGCGGTTGGGAATGTAATTCAGCTCCGCCATCGCCGCTTCCACTTTTTCCCGCGTTTTCGCAGA
AACGTGGCTGGCCTGGTTCACCACGCGGGAAACGGTCTGATAAGAGACACCGGCATACTCTGCGAC
ATCGTATAACGTTACTGGTTTCACATTCACCACCCTGAATTGACTCTCTTCCGGGCGCTATCATGCCAT
ACCGCGAAAGGTTTTGCGCCATTCGATGGTGTCCGGGATCTCGACGCTCTCCCTTATGCGACTCCTGC
ATTAGGAAGCAGCCCAGTAGTAGGTTGAGGCCGTTGAGCACCGCCGCCGCAAGGAATGGTGCATGC
AAGGAGATGGCGCCCAACAGTCCCCCGGCCACGGGGCCTGCCACCATACCCACGCCGAAACAAGCG
CTCATGAGCCCGAAGTGGCGAGCCCGATCTTCCCCATCGGTGATGTCGGCGATATAGGCGCCAGCAA
CCGCACCTGTGGCGCCGGTGATGCCGGCCACGATGCGTCCGGCGTAGAGGATCGAGATCTCGATCC
CGCGAAATTAATACGACTCACTATAGGGGAATTGTGAGCGGATAACAATTCCCCTCTAGAAATAATT
TTGTTTAACTTTAAGAAGGAGATATACCATGAAAAAGACAGCTATCGCGATTGCAGTGGCACTGGCT
GGTTTCGCTACCGTAGCGCAGGCCGCTCCGCAAGATAACACTAGCGCCATGGGCAAACATCACCATC
ACCATCACCCCATGAGCGATTACGACATCCCCACTACTAAGCTTCTGGAAGTTCTGTTCCAGGGGCCC
CATATGACTGGCTGTGTATAAGGGAGCCTGACATTTATATTCCCCAGAACATCAGGTTAATGGCGTTT
TTGATGTCATTTTCGCGGTGGCTGAGATCAGCCACTTCTTCCCCGATAACGGAGACCGGCACACTGG
CCATATCGGTGGTCATCATGCGCCAGCTTTCATCCCCGATATGCACCACCGGGTAAAGTTCACGGGA
GACTTTATCTGACAGCAGACGTGCACTGGCCAGGGGGATCACCATCCGTCGCCCGGGCGTGTCAATA

ATATCACTCTGTACATCCACAAACAGACGATAACGGCTCTCTCTTTTATAGGTGTAAACCTTAAACTGC
ATTTCACCAGCCCCTGTTCTCGTCAGCAAAAGAGCCGTTCATTTCAATAAACCGGGCGACCTCAGCCA
TCCCTTCCTGATTTTCCGCTTTCCAGCGTTCGGCACGCAGACGACGGGCTTCATTCTGCATGGTTGTG
CTTACCAGACCGGAGATATTGACATCATATATGCCTTGAGCAACTGATAGCTGTCGCTGTCAACTGTC
ACTGTAATACGCTGCTTCATAGCATACCTCTTTTTGACATACTTCGGGTATACATATCAGTATATATTC
TTATACCGCAAAAATCAGCGCGCAAATACGCATACTGTTATCTGGCTTTTAGTAAGCCGGATCCACGC
GTCTCGAGCACCACCACCACCACCACTGAGATCCGGCTGCTAACAAAGCCCGAAAGGAAGCTGAGTT
GGCTGCTGCCACCGCTGAGCAATAACTAGCATAACCCCTTGGGGCCTCTAAACGGGTCTTGAGGGGT
TTTTTGCTGAAAGGAGGAACTATATCCGGAT
